# Age dependency and lateralization in the three branches of the human superior longitudinal fasciculus

**DOI:** 10.1101/2021.01.27.428023

**Authors:** Kaoru Amemiya, Eiichi Naito, Hiromasa Takemura

**Affiliations:** Center for Information and Neural Networks (CiNet), National Institute of Information and Communications Technology, and Osaka University, Suita, Japan; Graduate School of Frontier Biosciences, Osaka University, Suita, Japan

**Keywords:** superior longitudinal fasciculus, diffusion MRI, quantitative MRI, age dependency, lateralization

## Abstract

The superior longitudinal fascicle/fasciculus (SLF) is a major white matter tract connecting the frontal and parietal cortices in humans. Although the SLF has often been analyzed as a single entity, several studies have reported that the SLF is segregated into three distinct branches (SLF I, II, and III). They have also reported the right lateralization of the SLF III volume and discussed its relationship with lateralized cortical functions in the fronto-parietal network. However, to date, the homogeneity or heterogeneity of the age dependency and lateralization properties of SLF branches have not been fully clarified. Through this study, we aimed to clarify the age dependency and lateralization of SLF I-III by analyzing diffusion-weighted MRI (dMRI) and quantitative R1 (qR1) map datasets collected from a wide range of age groups, mostly comprising right-handed children, adolescents, adults, and seniors (6 to 81 years old). The age dependency in dMRI measurement (fractional anisotropy, FA) was heterogeneous among the three SLF branches, suggesting that these branches are regulated by distinct developmental and aging processes. Lateralization analysis on SLF branches revealed that the right SLF III was larger than the left SLF III in adults, replicating previous reports. FA measurement also suggested that, in addition to SLF III, SLF II was lateralized to the right hemisphere in adolescents and adults. We further found a left lateralization of SLF I in qR1 data, a microstructural measurement sensitive to myelin levels, in adults. These findings suggest that the SLF sub-bundles are distinct entities in terms of age dependency and lateralization.

## 1. Introduction

Human brain comprises a number of functional systems to mediate various types of information processing and support behavior in daily life. The fronto-parietal network, a group of regions in the lateral prefrontal and posterior parietal cortices, is one of the prominent functional systems involved with essential cortical functions, such as visuospatial attention (Corbetta & Shulman, 2002), working memory (Linden et al., 2003; Rottschy et al., 2012), generation of motor intention and awareness (Desmurget et al., 2009), and adaptive cognitive control (Cole et al., 2013). Recently, functional MRI (fMRI), cortical stimulation, and neuropsychological studies have provided converging evidence that the functional organization of the human fronto-parietal network is not symmetric across hemispheres. For example, previous studies have demonstrated that the fronto-parietal network in the right hemisphere is taking a dominant role in neural processing on attention/spatial awareness (Corbetta et al., 2005; Shinoura et al., 2009; Thiebaut de Schotten et al., 2005), self-face recognition (Morita et al., 2018, 2020) or proprioceptive/motor awareness of limb movement (Amemiya & Naito, 2016; Berlucchi & Aglioti, 1997; Cignetti et al., 2014; Daprati et al., 2010; Naito et al., 2016).

White matter tracts connecting distant brain regions are essential for understanding the functional organization of cortical networks (Catani & Thiebaut de Schotten, 2012; Takemura & Thiebaut de Schotten, 2020). While the importance of white matter tracts has been documented by classical neuroanatomists (Catani & Ffytche, 2005), recent progress in diffusion-weighted MRI (dMRI) and tractography have revealed a more direct relationship between white matter tracts and behavioral data (Thiebaut de Schotten et al., 2011; Yeatman et al., 2012a). Neurobiological studies have provided evidence on the mechanisms underlying developmental changes in and the plasticity of white matter tracts, by revealing oligodendrocytes’ role on myelin plasticity and that such plasticity affects behavioral properties (Fields, 2015; Makinodan et al., 2012; Sampaio-Baptista et al., 2013; Wake et al., 2015). Therefore, converging evidence across research fields shows the importance of understanding the properties of white matter tracts in order to understand cortical functions.

The superior longitudinal fasciculus/fascicle (SLF) is one of the major white matter tracts connecting the frontal and parietal cortices and is thus considered to carry information among cortical areas belonging to the fronto-parietal network (Déjerine, 1895; Mori & van Zijl, 2002; Schmahmann & Pandya, 2006; Thiebaut de Schotten et al., 2011). Although a number of human dMRI studies have analyzed the human SLF as a single entity (De Santis et al., 2016; Hoeft et al., 2007; Lebel et al., 2012; Wakana et al., 2007; Yeatman et al., 2014), tracer studies in macaque (Petrides & Pandya, 1984; Schmahmann & Pandya, 2006) as well as dissection studies in humans (Komaitis et al., 2019; Thiebaut de Schotten et al., 2011), have provided evidence that the SLF has three branches (SLF I, II, and III), each of which terminates in different parts of the parietal and frontal cortices. Based on these anatomical studies, dMRI studies have successfully identified three SLF branches in living humans using tractography (Kamali et al., 2014; Makris et al., 2005; Schurr et al., 2019b; Thiebaut de Schotten et al., 2011, 2012; Wang et al., 2016). Further comparisons of dMRI-based measurements on SLF branches with functional or behavioral data have suggested that these branches may be involved in different types of functions (Budisavljevic et al., 2017; Cazzoli & Chechlacz, 2017; Howells et al., 2018; Parlatini et al., 2017; Thiebaut de Schotten et al., 2011; see Discussion). Therefore, it is crucial to investigate the structural properties of SLF branches in order to understand the cortical functions mediated by interactions among the respective cortical areas connected by SLF I, II, and III.

Several dMRI studies have revealed that the human SLF III in the right hemisphere has a larger volume than that in the left hemisphere (Budisavljevic et al., 2017; Hecht et al., 2015; Thiebaut de Schotten et al., 2011), suggesting a possible link between SLF III and right-dominant function in the fronto-parietal network (Amemiya & Naito, 2016; Cazzoli & Chechlacz, 2017; Chechlacz et al., 2015a, 2015b; Howells et al., 2018; Morita et al., 2018; Naito et al., 2016). Since some of these right-lateralized functions emerge during developmental processes (Morita et al., 2018; Naito et al., 2017), it is important to understand the details of age dependency and lateralization of the three SLF branches. Moreover, it is not yet fully understood whether these three branches should be considered as a single entity or as distinct components in terms of age dependency and lateralization.

The present study aimed to address two fundamental questions regarding the human SLF by analyzing a dMRI dataset collected from a large population of variable ages (from 6 to 81 years old; Yeatman et al., 2014). First, while the age dependency of white matter tracts has been assessed in previous dMRI studies (Lebel et al., 2012; Peters et al., 2014; Yeatman et al., 2014), the specific maturation and aging processes of the three branches of the human SLF have not been examined. Therefore, we examined the similarities or differences in age dependency among three branches, by analyzing the dMRI dataset collected from a wide range of age groups and subdividing the SLF into three distinct branches. Second, we evaluated the lateralization of SLF branches to establish how the properties of each branch may be associated with the functional lateralization of different parts of the fronto-parietal network. Although this analysis has been previously performed (Budisavljevic et al., 2017; Hecht et al., 2015; Thiebaut de Schotten et al., 2011), we aimed to further test SLF lateralization in participants of various ages.

For addressing each question, we analyzed not only macroscopic measurement (tract volume) but also two different types of microstructural measurements of the three SLF branches. Specifically, we analyzed fractional anisotropy (FA; Basser & Pierpaoli, 1996), a diffusion tensor model-based metric widely used for quantifying white matter microstructural properties. However, it is known that FA is not a specific measurement correlating directly with a single neurobiological factor (Assaf et al., 2019; Jones et al., 2013; Rokem et al., 2017; Sampaio-Baptista & Johansen-Berg, 2017; Thomason & Thompson, 2011; Wandell & Le, 2017), and thus, neurobiological interpretation of FA results remains ambiguous. In addition to FA, we also analyzed quantitative R1 (qR1) data included in the same dataset. Compared with FA, qR1, which is the inverse of quantitative spin-lattice relaxation time T1, is relatively specific to the myelin volume fraction, and thus provides insights on underlying neurobiological mechanisms explaining tissue properties along white matter tracts (Schurr et al., 2018, 2019a; Stüber et al., 2014; Takemura et al., 2019; Yeatman et al., 2014). Nevertheless, qR1 is not a fully specific measurement of myelin (Harkins et al., 2016; see Discussion). Therefore, we combined FA and qR1 data to elucidate the neurobiological basis of age dependency and lateralization of the human SLF system.

## 2. Material and Methods

### 2.1. MRI datasets

We analyzed the dMRI and qR1 dataset collected on a 3T General Electric Discovery 750 (General Electric Healthcare, Milwaukee, WI, USA) equipped with a 32-channel head coil (Nova Medical, Wilmington, MA, USA) at the Center for Cognitive and Neurobiological Imaging at Stanford University (www.cni.stanford.edu). This dataset has already been analyzed in previous publications (Bain et al., 2019a, 2019b; Erramuzpe et al., 2021; Lerma-Usabiaga et al., 2020; Schurr et al., 2019b; Yeatman et al., 2014). Data collection procedures were approved by the Stanford University Institutional Review Board. Participants were recruited from the San Francisco area and were screened for neurological, cognitive, and psychiatric disorders. Participants were not screened for disorders likely to occur later in life, such as hypertension. This dataset did not include the score of cognitive tests evaluating aging. Moreover, it purposefully included more children and adolescents, since participants of these ages were expected to show the largest age-dependency in white matter properties. All participants provided written informed consent, which was conducted in accordance with the ethical standards stated in the Declaration of Helsinki. Further details of the data acquisition have been described in a previous publication (Yeatman et al., 2014). Notably, no part of the study procedures and analyses had been pre-registered before conducting the research.

From the original 102 participants, we excluded participants who had not completed the high angular resolution dMRI scan. We analyzed the dMRI and qR1 dataset of 89 healthy participants of variable ages (ranged from 6 to 81 years old). We removed the data of seven participants who exhibited excessive motion during dMRI data acquisition (> 2.5 voxels in translation or > 1 degree in rotation on average), and divided the remaining 82 participants into four age groups: child (n = 17, range = 6-9 and average = 8.35 years old, male = 10), adolescent (n = 20, range = 10-18, average = 13.45 years old, male = 10), adult (n = 23, range = 20-50, average = 32.48 years old, male = 11), and senior (n = 22, range = 55-81, average = 64.86 years old, male = 10), for subsequent analyses. The majority of participants included in the analysis were right-handers (16 children, 16 adolescents, 15 adults, and 19 seniors). Only a few participants were reported to be left-handers (2 adolescents, 1 adult, and 1 senior). For the remaining participants, handedness had not been recorded.

In addition to the main group analysis mentioned above, we performed supplementary analysis using different group definitions: 1) a subgroup of adolescents only including participants aged between 12 and 18 years (n = 14; average = 14.64 years old, male = 7), 2) two subgroups of seniors, one including participants aged between 55 and 62 years (n = 11; average = 58.45 years old, male = 5) and the other including participants between 64 and 81 years old (n = 11; average = 71.27 years old; male = 5).

### 2.2. dMRI data acquisition and analyses

#### 2.2.1. dMRI data acquisition

The dMRI dataset was acquired using dual-spin echo diffusion-weighted sequences with full brain coverage (Reese et al., 2003) and 2 mm isotropic voxels. Acquisition included diffusion-weighted (b = 2000 s/mm^2^; 96 directions) and eight non-diffusion weighted (b = 0 s/mm^2^) images. The repetition time was 7,800 ms and the echo time was 93.7 ms.

#### 2.2.2. dMRI data preprocessing

Eddy current distortions and participant motion in dMRI images were removed by a 14-parameter, constrained non-linear co-registration based on the expected pattern of eddy-current distortions, given the phase-encode direction of the acquired data (Rohde et al., 2004) using mrDiffusion tools implemented in vistasoft distribution (https://github.com/vistalab/vistasoft). Diffusion gradients were adjusted to account for the rotation applied to the measurements during motion correction. The dMRI data was then coregistered with synthetic T1-weighted images obtained from the qR1 dataset (see below), which was aligned to the AC-PC (Anterior Commissure-Posterior Commissure) space. The diffusion tensor model was fitted to each voxel’s data by using the least-squares algorithm to calculate FA in each voxel. The FA was used to quantify microstructural properties along the SLF. The constrained spherical devolution (CSD; *L*_*max*_ = 6; Tournier et al., 2007) model was also fitted with each voxel’s data using MRTrix 3.0 (http://www.mrtrix.org/; Tournier et al., 2012, 2019) to estimate the fiber orientation distribution for performing CSD-based tractography.

#### 2.2.3. Fiber tracking

For each participant and each dataset, we used CSD-based deterministic tractography implemented in MRTrix 3.0 (SD_STREAM; Tournier et al., 2012, 2019) to generate 2 million streamlines (tractogram) for each dMRI dataset (step size, 0.2 mm; maximum angle between successive steps, 45°; minimum length, 10 mm; maximum length, 250 mm; fiber orientation distribution amplitude stopping criterion, 0.1). The angle threshold was identical to that used in a previous work reporting the SLF laterality (Thiebaut de Schotten et al., 2011). We used the entire white matter mask as seed, and seed voxels were randomly chosen from the mask for generating individual streamlines. Tracking was terminated when a streamline reached outside the white matter mask.

#### 2.2.4. Tract identification

##### Main analysis

From each tractogram, we identified the three branches of the SLF in all hemispheres using the multiple waypoint regions of interest (ROIs) approach proposed in previous studies (Rojkova et al., 2016; Thiebaut de Schotten et al., 2011). For each participant, we manually delineated all ROIs on synthetic T1-weighted images obtained from the qR1 dataset (see below), which was coregistered with the dMRI dataset. First, we manually defined three distinct frontal coronal ROIs for an “AND” operation at a coronal slice including the anterior commissure. Each ROI covered the superior frontal gyrus, middle frontal gyrus or precentral gyrus and was used for identifying SLF I, II, and III, respectively (cyan, blue, and magenta; Supplementary Figure 1A). These ROIs did not cover the white matter near the cingulate gyrus to exclude the cingulum bundle. Second, we delineated another coronal “AND” ROI that covered the parietal white matter regions superior to the lateral fissure, in a posterior coronal section including the posterior commissure (green; Supplementary Figure 1B). Third, we delineated a “NOT” ROI in the mid-sagittal plane to exclude the callosal fibers (yellow; Supplementary Figure 1A and 1B). Finally, we defined a “NOT” ROI in the axial slice covering the temporal lobe in order to exclude the arcuate fasciculus (Supplementary Figure 1C-D). Supplementary Figure 1 depicts the positions of waypoint ROIs overlaid on a synthetic T1-weighted image in a representative participant.

SLF streamlines were refined by a subsequent process of outlier streamline removal. Specifically, we removed streamlines that met the following criteria: (1) streamline length ≥ 4 standard deviation (SD) longer than the median streamline length in the tract, (2) streamline position ≥ 4 SD away from the median position of the tract (Yeatman et al., 2012b). Figure 1 depicts SLF I, II and III identified through these procedures in a representative participant from each group, visualized by using the MATLAB Brain Anatomy toolbox (https://github.com/francopestilli/mba).

##### Supplementary analysis using exclusive ROIs

We also performed a supplementary analysis using more exclusive ROI-based tract identification criteria for SLF I, II and III. This analysis aimed to evaluate he generalizability of the results of the main analysis when reducing the spatial overlaps among SLF I, II and III. To this end, in addition to the ROI-based SLF identification procedures used in the main analysis, we excluded the following streamlines: (1) SLF I streamlines passing through ROIS of either the middle frontal gyrus or precentral gyrus, (2) SLF II streamlines passing through ROIs of either the superior frontal gyrus or precentral gyrus, and (3) SLF III streamlines passing through ROIs of either the superior frontal gyrus or middle frontal gyrus. SLF streamlines were refined using the identical outlier removal criteria, as used in the main analysis.

### 2.3. qR1 data acquisition and analyses

The qR1 data were acquired for all participants following protocols described in previous publications (Gomez et al., 2017; Mezer et al., 2013; Yeatman et al., 2014). Four spoiled gradient echo (SPGR) images were acquired with flip angles of 4°, 10°, 20°, and 30° (repetition time, 14 ms; echo time, 2.4 ms), and a scan resolution of 1 mm isotropic. In addition, four additional spin echo inversion recovery (SEIR) images were acquired with an echo planar imaging (EPI) readout (repetition time, 3000 ms; echo time was set to minimum full) to remove field inhomogeneity. The inversion times were 50, 400, 1200, and 2400 ms. In-plane resolution and slice thickness of the additional SEIR images were 2 × 2 mm^2^ and 4 mm, respectively. Further details on qR1 data acquisitions in this dataset have been described in previous publications (Bain et al., 2019b; Yeatman et al., 2014).

Both the SPGR and SEIR images were processed using the mrQ software package (https://github.com/mezera/mrQ) in MATLAB to produce the quantitative T1 maps (Mezer et al., 2013). The mrQ analysis pipeline corrects for RF coil bias using SEIR-EPI scans (Barral et al., 2010), producing accurate T1 fits across the brain. We then calculated quantitative R1 (qR1) maps by calculating an inverse of quantitative T1 in each voxel. The full analysis pipeline and its published description can be found at https://github.com/mezera/mrQ (Mezer et al., 2013, 2016).

The mrQ analysis pipeline generates a T1-weighted image (synthetic T1-weighted image) from SPGR and SEIR images (Bain et al., 2019b). This synthetic T1-weighted image was used for co-registering dMRI data into the qR1 data space and for data visualization (Figure 1).

**Figure 1.**
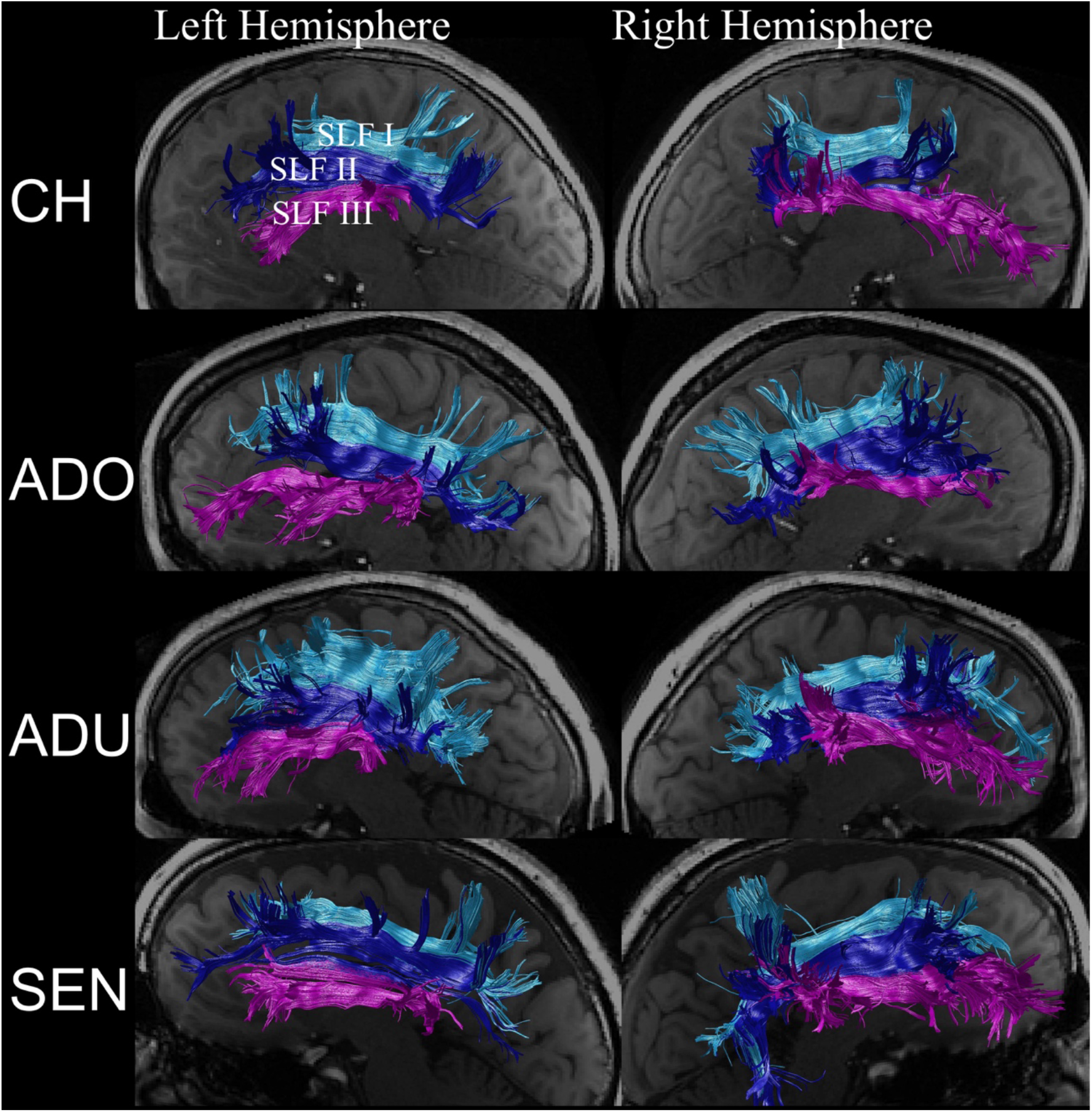
Three SLF branches (cyan, SLF I; dark blue, SLF II; purple, SLF III) identified from the dMRI dataset collected from a representative participant of each group (P1 from child; P18 from adolescent; P38 from adult; P61 from senior). The tracts in each hemisphere (left panels, left hemisphere; right panels, right hemispheres) are overlaid on a sagittal slice of a synthetic T1-weighted image obtained from the qR1 dataset located medial to the tracts. CH, child; ADO, adolescent; ADU, adult; SEN, senior; SLF, superior longitudinal fasciculus/fascicle.

### 2.4. Quantification and statistical analysis

#### 2.4.1. Tract volume

For each participant, we calculated the estimated volumes of SLF I, II, and III in each hemisphere by counting the number of voxels intersecting with single or multiple streamlines in the qR1 image (1 mm isotropic). We also performed a supplementary analysis of the tract volume by only counting voxels intersecting more than a certain number (5, 10 or 20) of streamlines, to evaluate the generality of results across choices of streamline density thresholds.

#### 2.4.2. Microstructural properties along tracts

We evaluated the microstructural properties of SLF I, II, and III based on the methods used in previous studies (Duan et al., 2015; Levin et al., 2010; Ogawa et al., 2014; Takemura et al., 2019, 2020; Yeatman et al., 2012b). We resampled each streamline to 100 equidistant nodes. Microstructural properties (FA and qR1) were calculated at each node of each streamline and summarized by taking the weighted average of measurements on each streamline within that node. The weight of each streamline was based on Mahalanobis distance from the tract core, which was calculated as the mean of each streamline’s x, y, z coordinates at each node (Yeatman et al., 2012b). We excluded the first and last 20 nodes from the microstructural property of the tract core to exclude voxels close to gray/white matter interfaces, where the tract was likely to intersect heavily with other fibers, such as the superficial U-fiber system. We summarized the profile of each tract using a vector of 60 values representing the microstructural measurements sampled at equidistant locations along the central portion of the tract (see Supplementary Figure 2 for depicting the position of these 60 nodes in a representative participant). In order to measure the age dependency and lateralization of the SLF branches, we used microstructural measurements (FA and qR1) averaged across 60 nodes in each hemisphere in subsequent analyses.

#### 2.4.3. Evaluating age dependency

##### Statistical Analysis

We evaluated the statistical difference between six tracts (left/right SLF I, II, and III) and four age groups (child, adolescent, adult, and senior) for tract volume or microstructural properties (FA and qR1) by performing a two-way analysis of variance (ANOVA) for assessing the main effect of the tracts and age groups and the interaction between tracts and age groups in each measurement. For measurements with a significant main effect of tracts or age groups, we performed a post-hoc test comparing the pairs of tracts or pairs of age groups using Sidak’s method. For measurements showing a significant interaction between the tracts and age groups, we further performed a post-hoc, simple main effect analysis to evaluate the difference in measurements in each tract between all pairs of age groups. The statistical significance (*P* value) of this analysis was corrected for multiple comparisons using Sidak’s method. Statistical tests have been performed by using SPSS Statistics 26 (IBM, New York).

##### Curve fitting

We further evaluated the age dependency of SLF properties by fitting a model to explain tract properties as a function of the participant’s age. We used a Poisson curve as used in previous studies (Lebel et al., 2012; Yeatman et al., 2014) with the following equation:

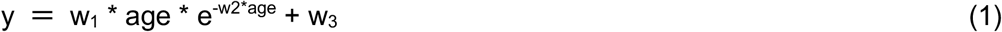

where w_1_, w_2_, and w_3_ are parameters estimated using the Levenberg–Marquardt algorithm with a least-squares cost function. We estimated the 95% confidence interval of the age of peak in age-dependency curves using the bootstrap method, as described in a previous study (Yeatman et al., 2014; https://github.com/jyeatman/lifespan).

##### Multiple linear regression

Since we found a significant interaction between age group and tracts in FA measurements and SLF I showed the largest age dependency (Figure 3), we performed a post-hoc analysis for evaluating how much of the age dependency of FA measurements along SLF I could be explained by variance in FA measurements along SLF II and III with a constant (*c*), by performing multiple linear regression analysis.

First, we fit the multiple linear regression model for predicting the variance in FA along SLF I based on FA variance along SLF II and III:

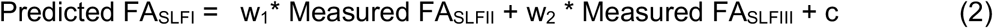

We performed this analysis by using the MATLAB Statistics and Machine Learning Toolbox aiming to minimize the least squared error by selecting the best combination of weights and constant.

We then calculated the residual FA along SLF I by calculating the difference between the measured and predicted FA values.

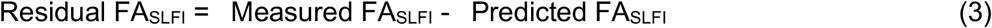

The residual FA of SLF I describes the inter-participant variance, which could not be explained by the variance in FA along SLF II and III. We evaluated the age dependency of the residual FA of SLF I for quantifying the extent to which it is independent from that of the FA of SLF II and III.

#### 2.4.4. Evaluating lateralization

##### Lateralization Index

For both tract volume and microstructural measurements (FA and qR1) along the tracts, we quantified the degree of lateralization by calculating the laterality index (LI) as used in previous studies (Bain et al., 2019b; Thiebaut de Schotten et al., 2011):

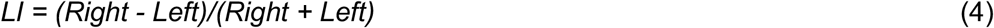

A positive LI indicated right lateralization whereas a negative LI indicated left lateralization. The LI was calculated separately for each age group.

In addition to the LI, we also calculated the effect size and statistical significance of left-right asymmetry in each tract (SLF I, II, and III) and metrics (volume, FA, and qR1) separately for each group (child, adolescent, adult, and senior). The effect size was estimated by calculating Cohen’s d’ on the difference between the left and right hemispheres. The statistical significance was assessed by the two-tailed paired t-test. For each test, we defined statistical significance (α) as *P* = 0.004, which is equivalent to *P* = 0.05 after Bonferroni correction for 12 comparisons (three tracts and four age groups).

#### 2.4.5. Comparison between female and male participants

For tract volume, FA, and qR1 in each tract and age group, we performed statistical comparisons between male and female participants using two-tailed two-sample t-test. For each test, we defined statistical significance (α) as *P* = 0.002, which is equivalent to *P* = 0.05 after Bonferroni correction for 24 comparisons (six tracts and four age groups).

#### 2.4.6. Evaluating tract overlap

We evaluated the spatial overlap among tracts (Kaneko et al., 2020; Sani et al., 2019) by calculating the proportion of voxels intersecting multiple SLF branches (SLF I, II, and III). We quantified the degree of spatial overlap by calculating Dice coefficient between a pair of SLF branches (SLF I/II, SLF II/III, and SLF I/III) in each hemisphere.

We also quantified the degree of overlap among SLF I, II, and III, identified by exclusive ROIs (see above), to evaluate the reduction in the spatial overlap as compared with that observed in the main analysis. In addition, we analyzed the tract volume, FA, and qR1 for SLF I, II, and III identified by exclusive ROIs using a procedure identical to that used in the main analysis, in order to evaluate how much the spatial overlap’s reduction affects the main findings.

## 3. Results

Using CSD-based tractography for dMRI data, we identified three branches of the SLF in all the 164 hemispheres analyzed (see Material and Methods for details). Figure 1 depicts the three SLF branches in a representative participant in each age group. In all hemispheres, SLF I, II, and III appeared as tracts connecting the parietal cortex with the superior frontal gyrus, middle frontal gyrus, and precentral/inferior frontal gyrus, respectively.

In subsequent analyses, we examined how much the SLF branches can be considered as a single entity or distinct system in terms of age dependency, microstructural property, and lateralization. We first investigated how much age dependency is homogeneous or heterogeneous across the three SLF branches. We evaluated age dependency of both macroscopic (tract volume) and microstructural (FA and qR1) properties of SLF I, II, and III. We then investigated the lateralization of SLF branches in terms of tract volume, in order to confirm right lateralization of SLF III as reported in previous studies. Furthermore, we also investigated the lateralization of FA and qR1 measurements in SLF I, II, and III, in order to improve our understanding on the neurobiological underpinnings of the lateralization of the human fronto-parietal network.

### 3.1. Age dependency of the macroscopic properties of the SLF

First, we evaluated the age dependency of the macroscopic property (tract volume) of the three SLF branches. To this end, we counted the number of voxels intersecting the SLF streamlines and quantified the volume of SLF branches in all individual hemispheres (see Material and Methods). Figure 2 depicts the SLF tract volume in each branch (SLF I, II, and III in the left and right hemispheres), averaged across hemispheres in each age group. We then statistically evaluated the tract volume data using two-way ANOVA for assessing the main effect of the age group and the tract.

**Figure 2.**
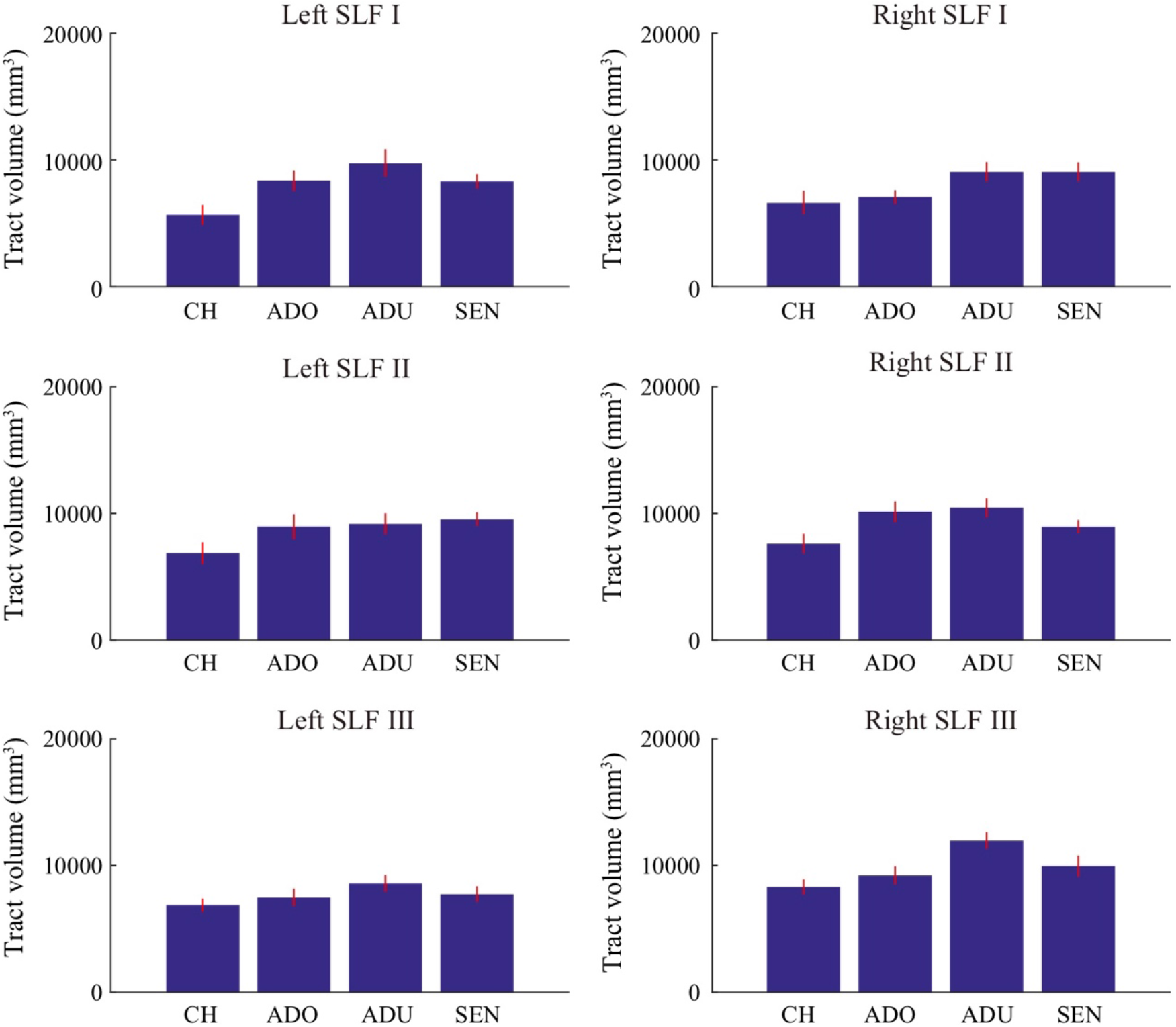
Estimated tract volume of the three SLF branches in the left and right hemisphere in each age group. The vertical axis depicts the average of tract volume across the hemispheres in each age group. The error bar depicts ±1 s.e.m. CH, child; ADO, adolescent; ADU, adult; SEN, senior; SLF, superior longitudinal fasciculus/fascicle.

We found a significant main effect of the age group (*F*_*3,468*_ = 16.20; *P* < 0.001). Post-hoc tests with Sidak’s method suggested statistically significant differences between children and all other age groups (adolescent, adult, and senior; *P* = 0.002, < 0.001, and < 0.001, respectively) and between adolescents and adults (*P* = 0.007). Other pairs did not show statistically significant differences (adolescent-senior and adult-senior; *P* = 0.91 and 0.11). Therefore, the significant main effect of the age group was mostly driven by a difference which may occurr during the developmental stage from children to adults.

We also found a main effect of the tract (*F*_*5,468*_ = 5.99; *P* < 0.001), indicating that tract volume was different across the SLF branches. In the post-hoc test, we found significant differences in some pairs (left SLF I - right SLF III, left SLF III - right SLF II, left SLF III - right SLF III, and right SLF I - right SLF III; P = 0.004, 0.01, < 0.001, and 0.002, respectively), but not in other pairs. A significant volume difference between the left and right SLF III suggested lateralization of this branch. Results of a comprehensive analysis of SLF III lateralization are described in section 3.3 (“Lateralization of tract volume in each age group”).

In contrast, we did not find a significant interaction between age group and tract (*F*_*15,468*_ = 0.90; *P* = 0.57), suggesting that while the tract volume of the SLF differs across age groups, there is no statistical evidence showing that such a maturation and aging profile is heterogeneous across SLF branches.

We further evaluated the age dependency of tract volume of the individual SLF branches by fitting a Poisson curve (Lebel et al., 2012; Yeatman et al., 2014; Supplementary Figure 3). However, since tract volume estimates had large inter-individual variability (Lebel et al., 2012), we could not obtain reliable estimates on the peak age of age-dependency curves.

In the main analysis, we counted the number of voxels intersecting with more than one SLF I/II/III streamlines for estimating the tract volume. We tested the degree to which age dependency of the tract volume depended on an arbitrary choice of streamline density threshold. To this end, we calculated tract volume by only counting voxels intersecting with a certain number (5, 10, and 20) of streamlines (see Material and Methods). While estimates of tract volume were reduced in more conservative streamline density thresholds, overall age dependency of the tract volume remained (Supplementary Figure 4). We also confirmed that the main effect of age group and tract remained significant at all thresholds (*P* < 0.001; Supplementary Table 1).

Taken together, these results provided profound evidence on the age dependency of the SLF volume, whereas there was no statistical evidence for supporting the heterogeneity of age dependency across SLF branches.

### 3.2. Age dependency of the microstructural properties of the SLF

We then evaluated the age dependency of the microstructural properties of the three SLF branches by two distinct measurements (FA and qR1) with different sensitivities for the underlying microstructural properties along the white matter tracts (Mezer et al., 2013; Takemura et al., 2019).

#### Age dependency of FA measurement

Figure 3 depicts the FA measurements along each tract and age group. FA measurements were averaged along the tract (see Material and Methods; see Supplementary Figure 5 for spatial profile). We analyzed FA measurements using two-way ANOVA in order to assess the main effect of age group and tract, and the interaction between them.

**Figure 3.**
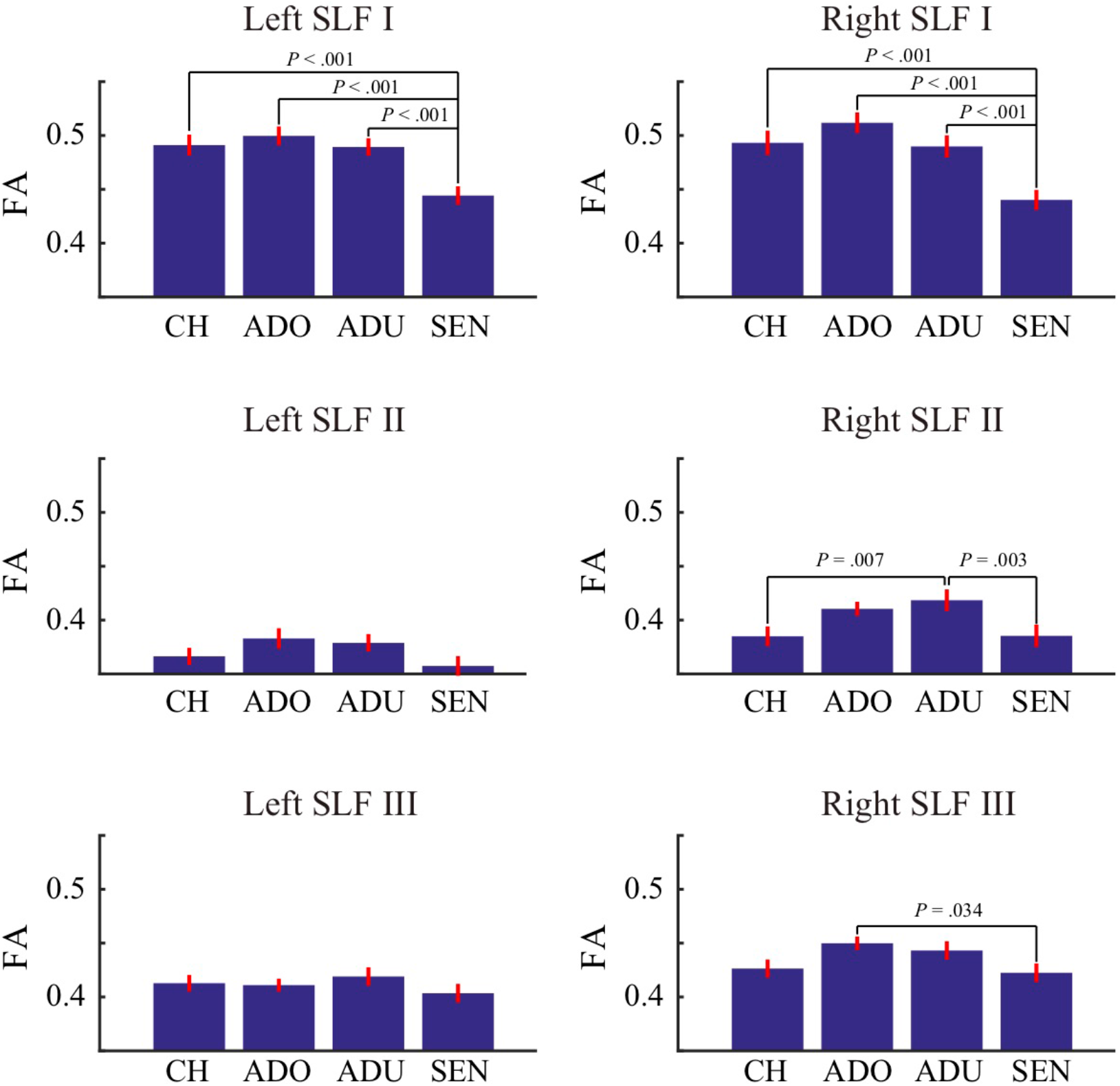
Estimated FA of the three SLF branches in the left and right hemisphere in each age group. The vertical axis depicts the average of FA across hemispheres in each age group. Statistically significant differences between a pair of age groups (post-hoc simple main effect analysis; *P* < 0.05 corrected by Sidak’s method) are also depicted in each plot. The error bar depicts ±1 s.e.m. CH, child; ADO, adolescent; ADU, adult; SEN, senior; SLF, superior longitudinal fasciculus/fascicle; FA, fractional anisotropy.

We found a significant main effect of the age group (*F*_*3,468*_ = 31.52; *P* < 0.001). Post-hoc analysis showed a significant difference between seniors and other groups (child, adolescent, and adult; *P* < 0.001 in all three cases) and between children and adolescents (*P* = 0.003). Other pairs did not show significant difference (child– adult and adolescent-adult; *P* = 0.07 and 0.84). This result suggests that the significant main effect of age group can largely be explained by decline of FA in seniors.

We also found a main effect of the tract (*F*_*5,468*_ = 161.51; *P* < 0.001), suggesting some microstructural differences across the three SLF branches. Post-hoc analysis showed significant differences in most pairs (*P* < 0.001 in all cases), except for the left SLF I/right SLF I (*P* = 1.00) and the left SLF III/right SLF II (*P* = 0.36). This result is consistent with that of a previous study showing that SLF I has the highest FA compared with that of the other two branches (Fitzgerald et al., 2018). These results also suggest, although the FA of SLF I is mostly symmetric across hemispheres, that of SLF II and III show significant differences between the left and right hemispheres, as discussed in section 3.3 (“Lateralization of tract volume in each age group”).

In addition to these main effects, we found a significant interaction between age group and tract (*F*_*15,468*_ = 2.84; *P* < 0.001). Since we found a significant interaction, we performed a post-hoc simple main effect analysis (see Material and Methods) to evaluate the significance of FA difference between each age group in each tract. For SLF I in both hemispheres, we found significant differences between seniors and the other three age groups (*P* < 0.001). For the right SLF II, we found significant differences between children and adults (*P* = 0.007) and between adults and seniors (*P* = 0.003). For the right SLF III, we found a significant difference between adolescents and seniors (*P* = 0.03). None of the differences between the other pairs of age groups reached statistical significance (*P* > 0.05, corrected by Sidak’s method). The significant interaction between age group and tract suggests that age dependency of FA measurements is heterogeneous across the three SLF branches.

We further assessed the age dependency of FA measurements along the SLF branches by fitting a Poisson curve (Figure 4). Curve fitting suggested that FA along SLF I in both hemispheres had clear decreasing trends during the aging process, though the peak of the curve could not be identified. FA measurements of SLF I may mature at early adolescent ages, but such a peak may be masked by large individual differences in children and adolescents. The right SLF II had a peak around early adult age (estimated peak age = 27; 95% CI of peak age, 21.6 - 33.8). In other tracts, we could not reliably identify the peaks of FA age-dependency curves.

**Figure 4.**
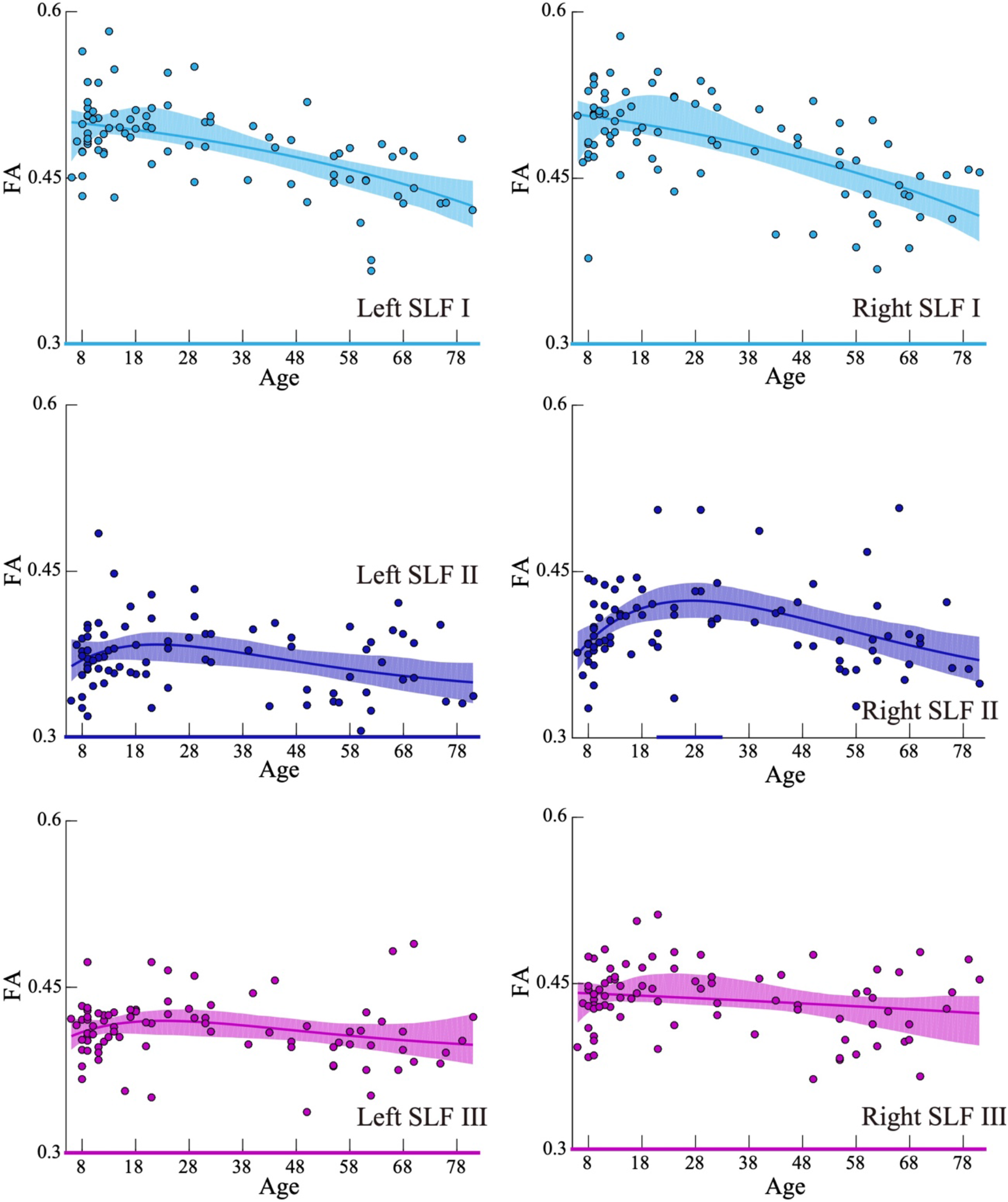
FA age-dependency curves for each SLF branch (left/right SLF I, II, and III). Each dot depicts data from an individual participant. The width of the curve denotes the 95% confidence interval around the Poisson curve fit. Colored lines at the bottom depicts the 95% confidence interval of the peak age, as estimated by the bootstrapping method (Yeatman et al., 2014). Except for the right SLF II, the reliable peak age of FA could not be identified. SLF, superior longitudinal fasciculus/fascicle; FA, fractional anisotropy.

Finally, we assessed the extent to which the age dependency of FA along SLF I was independent from that of FA along SLF II and III. To do so, we fitted a multiple linear regression model that predicts the FA of SLF I from the that of SLF II and III (see Material and Methods). This model predicted a modest amount of variance in the FA along SLF I (left hemisphere, R^2^ = 0.12; right hemisphere, R^2^ = 0.08). We then calculated the residual FA along SLF I, a variance that was not explained by that in FA of SLF II and III (see Material and Methods). Figure 5 depicts the age dependency of the residual FA along SLF I. Overall, we still observed decrement in the senior group in the residual FA along SLF I, suggesting that the age-dependency of the FA along SLF I cannot be fully explained by a variance in SLF II and III. This result suggests that, although some mechanisms underlying age dependency may be common across SLF branches, the age dependency profile of SLF I is distinct from that of SLF II and III, with differences being especially prominent at older ages.

**Figure 5.**
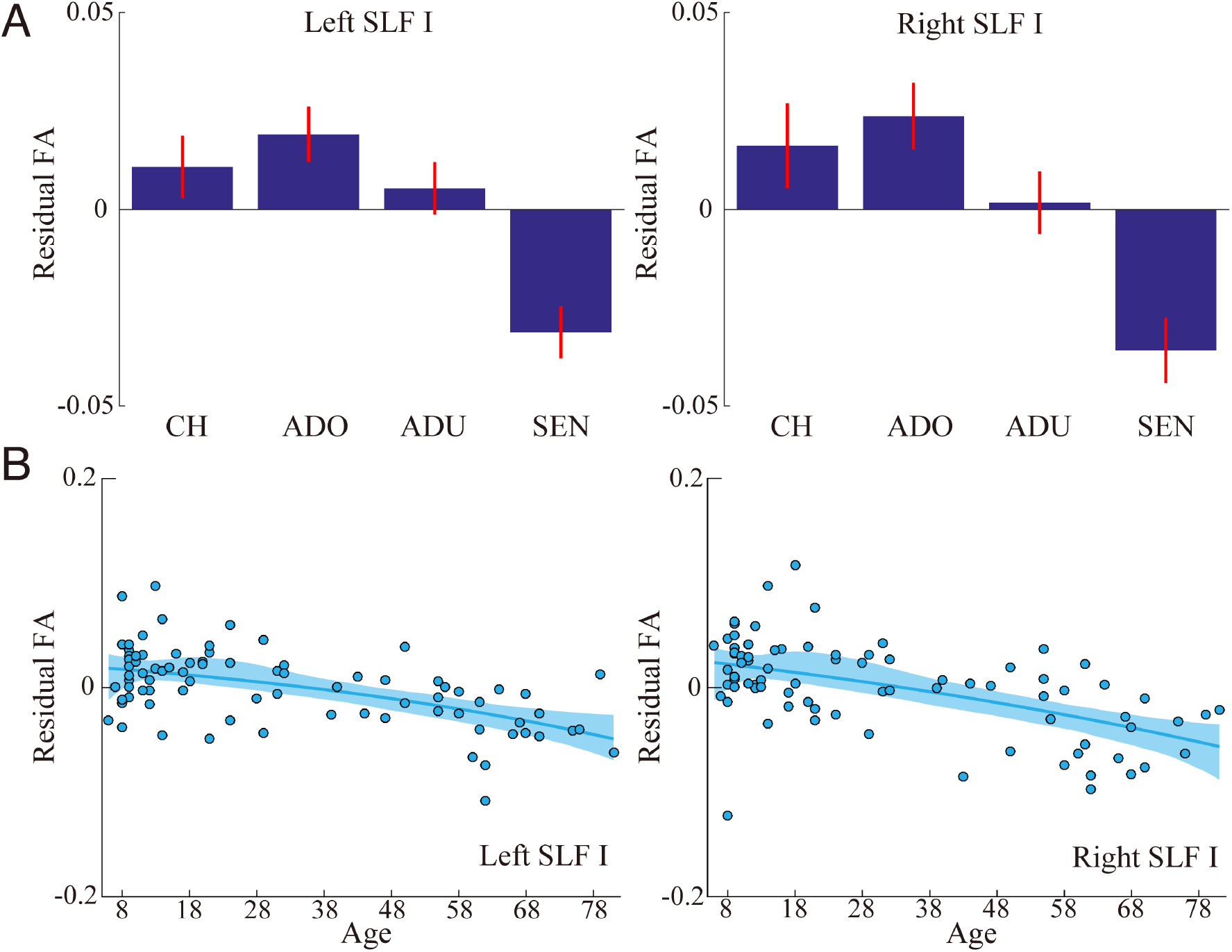
**A.** Comparison of residual FA on SLF I, an inter-participant variance that could not be explained by the variance of FA along SLF II and III (see Material and Methods), across all age groups (left panel, left SLF I; right panel, right SLF I). Conventions are identical to those in Figure 3. **B.** Age-dependency curve of the residual FA on SLF I (left panel, left SLF I; right panel, right SLF I). Conventions are identical to those in Figure 4. CH, child; ADO, adolescent; ADU, adult; SEN, senior; SLF, superior longitudinal fasciculus/fascicle; FA, fractional anisotropy.

#### Age dependency of qR1 measurement

Figure 6 depicts the qR1 measurements, which are sensitive to myelin levels, along each tract and age group (see Supplementary Figure 6 for spatial profile). We performed two-way ANOVA on qR1 measurements for assessing the statistical significance.

**Figure 6.**
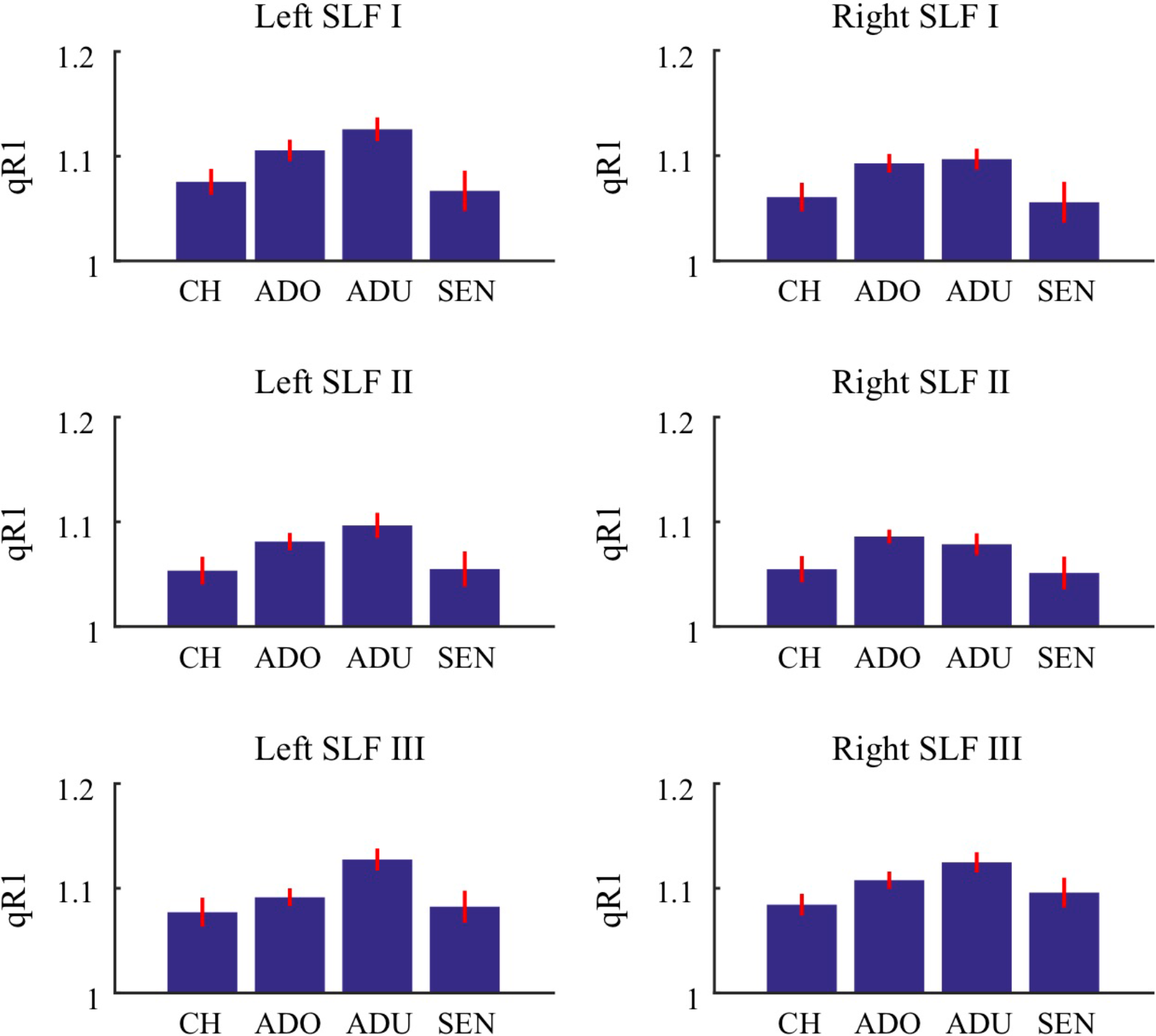
Estimated qR1 of the three branches of the SLF in the left and right hemisphere in each age group. The vertical axis depicts the average qR1 across hemispheres in each age group. The error bar depicts ±1 s.e.m. CH, child; ADO, adolescent; ADU, adult; SEN, senior; SLF, superior longitudinal fasciculus/fascicle; qR1, quantitative R1.

We found a significant main effect of the age group (*F*_*3,468*_ = 20.70; *P* < 0.001). Post-hoc analysis showed a significant difference between children and adolescents/adults (*P* = 0.001 and < 0.001, respectively) and between adolescents/adults and seniors (*P* < 0.001 in both cases). This result suggests that the age dependency of qR1 data may include both increasing trend during development and decreasing trend during aging.

We also found a main effect of the tract (*F*_*5,468*_ = 6.85; *P* < 0.001). Post-hoc analysis showed significant differences in some pairs of tracts (left SLF I – right SLF II, left SLF II – left SLF III, left SLF II – right SLF III, left SLF III – right SLF II, right SLF I – right SLF III, and right SLF II – right SLF III; *P* = 0.01, 0.04, 0.001, 0.005, 0.008, and < 0.001, respectively), but not in other pairs.

These results on qR1 measurements also suggest microstructural differences across SLF branches and age groups. However, we did not find a significant interaction between age group and tract for qR1 measurements (*F*_*15,468*_ = 0.52; *P* = 0.93), unlike that for FA measurements. Therefore, there is no statistical evidence supporting a heterogeneous age dependency of qR1 measurements across SLF branches.

Finally, we evaluated the age dependency of these tracts by fitting the Poisson curve (Supplementary Figure 7). In comparison with FA, the qR1 of the left SLF I clearly peaked at early adulthood (estimated peak age = 26; 95% CI of peak age, 21.4 - 30.9). The age-dependency curve was almost symmetric between the left and right SLF III, with an estimated peak at the age of 30 years in both hemispheres (95% CI of peak age, left SLF III, 24.1 - 36.5; right SLF III, 24.5 - 39.2). For the other tracts, we did not find a reliable peak in the age-dependency curve of qR1.

### 3.3. Lateralization of the tract volume in each age group

We investigated the degree of lateralization of the SLF I, II, and III volume in each age group by assessing the lateralization index (LI) and evaluating the statistical difference between the left and right hemisphere (see Material and Methods).

#### Lateralization of tract volume

The left panel in Figure 7 depicts the LI of the tract volume (see Supplementary Figure 8 for scatter plots comparing left and right hemispheres). For SLF I and II, we did not find statistically significant differences in volume across hemispheres in all age groups (SLF I: d’ = 0.32, −0.41, −0.22, and 0.18; *P* = 0.20, 0.09, 0.31, and 0.40; SLF II: d’ = 0.20, 0.29, 0.30, and −0.17; *P* = 0.42, 0.21, 0.16, and 0.43 for child, adolescent, adult, and senior, respectively). For SLF III, tract volume in the right hemisphere was significantly larger than that in the left hemisphere in adults (d’ = 1.32, *P* = 0.000002). A similar right lateralization of SLF III volume was observed in other age groups (child: d’ = 0.58, *P* = 0.03; adolescent: d’ = 0.49, *P* = 0.04; senior: d’ = 0.56, *P* = 0.02), although these effects did not reach statistical significance after Bonferroni correction (α = 0.004).

**Figure 7.**
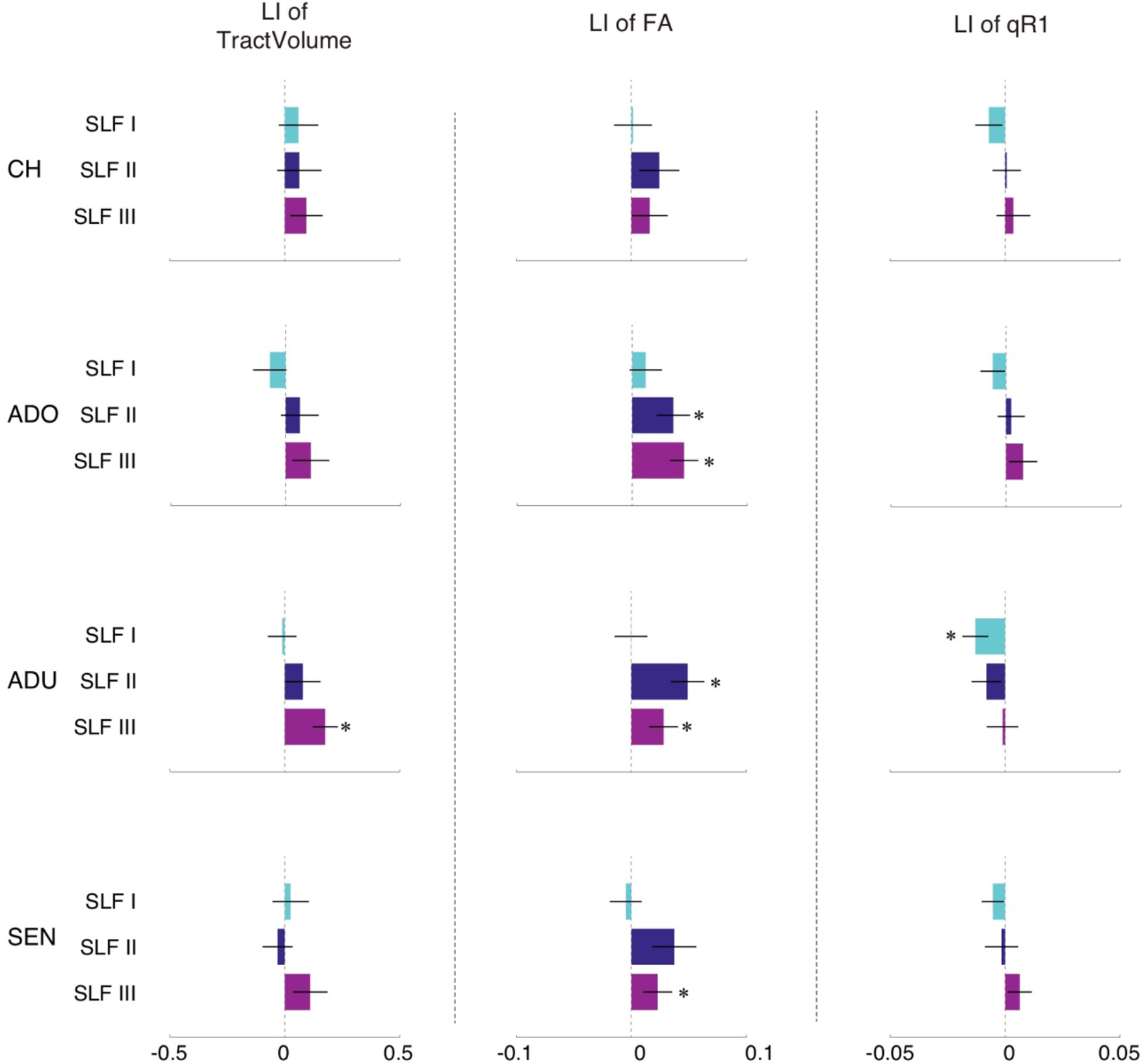
Lateralization index (LI). Hemispheric lateralization of each SLF branch (SLF I, II, and III) in each age group (child, adolescent, adult, and senior) assessed by the LI. A positive LI value indicates that the measurement along the tract is right lateralized. *Left panel*: LI of tract volume. *Middle panel*: LI of FA. *Right panel*: LI of qR1. Asterisks indicate statistically significant differences between the left and right hemispheres (*P* < 0.004). The error bar indicates ±1 s.e.m. across all participants. CH, child; ADO, adolescent; ADU, adult; SEN, senior; SLF, superior longitudinal fasciculus/fascicle; FA, fractional anisotropy; qR1, quantitative R1.

We also confirmed that the right lateralization of SLF III volume was maintained or was even increased when we varied the streamline density threshold (Supplementary Figure 9; see Material and Methods), indicating a generalization of lateralization across threshold choices.

Taken together, these results replicate the right lateralization of the tract volume of human SLF III in adults, as reported in previous studies (Budisavljevic et al., 2017; Hecht et al., 2015; Thiebaut de Schotten et al., 2011).

### 3.4. Lateralization of the microstructural property in each age group

Next, we assessed the degree of lateralization of the microstructural properties (FA and qR1) of SLF I, II, and III in each age group.

#### Lateralization of FA

The middle panel of Figure 7 depicts the LI of FA measurements along SLF I, II, and III in each age group (see Supplementary Figure 10A for the spatial profile, Supplementary Figure 11 for scatter plots). There was no significant lateralization of the FA of SLF I in any age group (d’ = 0.05, 0.31, 0.01, and −0.12; *P* = 0.84, 0.18, 0.95, and 0.57 for child, adolescent, adult, and senior, respectively). In contrast, we found a significant right lateralization of the FA of SLF II in adolescents (d’ = 0.87, *P* = 0.001) and adults (d’ = 1.15, *P* = 0.00002). SLF II in children and seniors also showed a similar right lateralization but the effect did not reach statistical significance after Bonferroni correction (d’ = 0.52 and 0.57; *P* = 0.05 and 0.01 for child and senior). Similar to SLF II, we also found a significant right lateralization of the FA of SLF III from adolescents to seniors (d’ = 1.54, 0.88, and 0.71; *P* = 0.000002, 0.0004, and 0.003 for adolescent, adult, and senior). No significant lateralization of the FA of SLF III was found in children (d’ = 0.39; *P* = 0.12). The spatial profile of the LI suggests that SLF III shows a greater right lateralization in its anterior part, presumably because its posterior part crosses with other tracts, such as the arcuate fasciculus (Supplementary Figure 10A). The above data provide profound evidence of the right lateralization of FA measurements not only for SLF III, but also for SLF II.

#### Lateralization of qR1

The right panel of Figure 7 depicts the LI of qR1 measurements (see Supplementary Figure 10B for the spatial profile, Supplementary Figure 12 for scatter plots). In contrast to the lack of lateralization of the tract volume and FA, we found significant left lateralization of qR1 of SLF I in adults (d’ = −0.98, *P* = 0.0001). SLF I also showed a similar left lateralization in other age groups, although the effect did not reach statistical significance after Bonferroni correction (d’ = −0.57, −0.54, and −0.61; *P* = 0.03, 0.03, and 0.01 for child, adolescent, and senior, respectively). We did not find significant evidence regarding lateralization of qR1 of SLF II and III in any age group, but we noted a modest left lateralization of SLF II in adults and a right lateralization of SLF III in adolescents and seniors, although these results did not reach statistical significance after Bonferroni correction (SLF II, d’ = 0.05, 0.17, −0.49, and −0.09; *P* = 0.85, 0.46, 0.03 and 0.67; SLF III, d’ = 0.18, 0.52, −0.06, and 0.59; *P* = 0.46, 0.03, 0.76, and 0.01 for CH, ADO, ADU, and SEN, respectively).

### 3.5. Lateralization of the SLF and handedness

We performed a supplementary analysis of the LI by excluding participants who were left-handed or did not have a record of handedness. Although this exclusion limited the statistical power because of the smaller number of participants (n = 16, 16, 15, and 19 for child, adolescent, adult, and senior), we aimed to evaluate how much the observed lateralization depended on handedness, since a previous study reported a relationship between SLF lateralization and handedness (Howells et al., 2018).

Supplementary Figure 13 depicts the LI of the tract volume, FA, and qR1 in right-handed participants. The overall pattern of lateralization was well preserved compared with that of the main analysis (Figure 7). Right lateralization of the SLF III volume in adults remained statistically significant (d’ = 1.06, *P* = 0.001), while that of the FA of SLF II remained significant in adults (d’ = 0.97, *P* = 0.002) but not in adolescents (d’ = 0.73, *P* = 0.01), presumably due to the reduced statistical power. Right lateralization of the FA of SLF III remained significant in adolescents (d’ = 1.50, *P* = 0.00002), but not in adults (d’ = 0.86, *P* = 0.005) and seniors (d’ = 0.68; *P* = 0.008). The left lateralization of qR1 of SLF I also remained significant in this analysis (d’ = −0.96, *P* = 0.002).

### 3.6. Comparison between male and female participants

For each age group, we evaluated statistical differences in tract volume, FA, and qR1 between male and female participants using two-sample t-test (Supplementary Figures 14-16). We did not find significant differences between male and female participants (*P* < 0.002; Bonferroni corrected for 24 comparisons) except that the FA along the right SLF III in seniors was significantly larger in males than in females (*P* = 0.001; Supplementary Figure 15; see Supplementary Table 2 for details).

### 3.7. Across-participant correlation of laterality measurements

We assessed the inter-participant correlation between lateralization measurements (the LI in volume of the SLF III, FA of the SLF II, FA of the SLF III and qR1 of the SLF I) in order to evaluate whether each lateralization properties were independent or not. This analysis was performed by pooling participants across all age groups (*N* = 82). We did not find any significant correlation among LI measurements (Supplementary Table 3), while there was a marginally significant correlation between the LI of the SLF III volume and that of the SLF III FA (R = 0.20, *P* = 0.07; Supplementary Figure 17). This result indicates that although we found profound evidence on the macroscopic (tract volume) and microscopic (FA and qR1) lateralization of SLF branches, each property might develop independently.

### 3.8. Evaluating the impact of spatial overlap between SLF I, II, and III

We evaluated the spatial overlap among the three SLF branches by quantifying the proportion of overlapping voxels using Dice coefficient in each pair of branches and age groups (blue bars, Supplementary Figure 18; see Material and Methods). There was almost no overlap between SLF I and III in all cases (Dice coefficient < 0.003). In contrast, SLF I and II shared 10-20% of voxels (left hemisphere: Dice coefficient = 0.15, 0.15, 0.12, and 0.20; right hemisphere: Dice coefficient = 0.10, 0.14, 0.14, and 0.15 for child, adolescent, adult, and senior, respectively). Similarly, SLF II and III also shared 7-15% of voxels (left hemisphere: Dice coefficient = 0.10, 0.07, 0.08, and 0.09; right hemisphere: Dice coefficient = 0.15, 0.11, 0.13, and 0.14 for child, adolescent, adult, and senior, respectively).

We then performed a supplementary analysis to test whether the results of the main analysis can be generalized upon the removal of voxels intersecting multiple SLF branches. To this end, we performed ROI-based identification of SLF I, II, and III in a more exclusive way: streamlines intersecting ROIs that were used to identify other branches were excluded from the analysis (“exclusive ROIs”, see Material and Methods). After these exclusions, the spatial overlap between SLF I/II and SLF II/III was substantially reduced (yellow bars, Supplementary Figure 18: left SLFI/II: 0.04, 0.05, 0.04, and 0.08; right SLF I/II: 0.03, 0.06, 0.06, and 0.06; left SLF II/III: 0.02, 0.02, 0.03 and 0.04; right SLF II/III: 0.06, 0.06, 0.04, and 0.07 for child, adolescent, adult, and senior, respectively). Using this SLF branch definition, we obtained results regarding the age group difference in tract volume (Supplementary Figure 19), FA (Supplementary Figure 20), and qR1 (Supplementary Figure 21), as well as regarding the lateralization (Supplementary Figure 22), that were consistent with those of the main analysis (see Supplementary Tables 4 and 5 for statistics on these analyses). Therefore, age group difference and lateralization results in the main analysis might not significantly depend on the existence of voxels intersecting multiple SLF branches.

### 3.9. Dependency on age group definitions

In the main analysis, we classified participants into four age groups (see Material and Methods). While these age groups were defined to approximately match the statistical power across groups, this definition did not account for the large age variability among participants in the adolescent and senior groups. To evaluate the dependency of age group definition, we tested two other age group definitions. First, we compared data of a subgroup of adolescents only including participants aged between 12 and 18 years, with those of the main analysis (including participants with 10-18 years old) but we did not find any notable differences (Supplementary Figure 23A). Second, we subdivided the senior group into two subgroups based on the participants’ age (*n* = 11 for each group, see Material and Methods for details). Again, we did not find any statistically significant differences between subgroups (Supplementary Figure 23B; tract volume: d’ = 0.15, −0.56, −0.23, −0.05, −0.31, and 0.17; *P* = 0.64, 0.09, 0.46, 0.88, 0.32, and 0.59; FA, d’ = −0.33, −0.56, −0.50, 0.05, −0.21, and −0.38; *P* = 0.30, 0.09, 0.13, 0.87, 0.51, and 0.24; qR1, d’ = 0.25, 0.39, 0.23, 0.19, 0.45, and 0.32; *P* = 0.42, 0.22, 0.46, 0.54, 0.17, and 0.31 for left SLF I, left SLF II, left SLF III, right SLF I, right SLF II, and right SLF III, respectively). While it was not possible to test all different types of age group definitions, we have prepared codes and data for replicating the analysis of this work that are publicly available (https://github.com/htakemur/SLFbranchesAgedependency), for encouraging readers to test their own age group definitions.

## 4. Discussion

In this work, we aimed to evaluate the age dependency of three SLF branches by analyzing dMRI and qR1 datasets collected from participants of various ages. We then evaluated the lateralization of the three branches in terms of measurements of tract volume and microstructural properties (FA and qR1).

### 4.1. Age dependency of the three SLF branches

A number of dMRI studies have investigated age dependency of the properties of white matter tracts, including the SLF (Kochunov et al., 2012; Lebel et al., 2012; Catherine Lebel & Beaulieu, 2011; Lynch et al., 2020; Slater et al., 2019; Westlye et al., 2010; Yeatman et al., 2014). These studies have revealed some heterogeneities in the maturation and aging processes across white matter tracts. However, they have not distinguished the three branches of the SLF. The current study reveals the age dependency of SLF I, II, and III separately.

We found a significant interaction between age group and SLF branches (SLF I, II, and III) in terms of FA measurements, strongly suggesting that the age dependency of diffusivity measurements is heterogeneous across SLF branches (Figures 3 and 4). This result suggests that, although the SLF has often been analyzed as a single white matter bundle, its branches are regulated by distinct developmental and aging processes.

Different age dependency profiles may be related to the functionality of these branches. For example, SLF I is considered to be associated with spatial and motor functions (Parlatini et al., 2017), while SLF III is considered to be associated with higher-order cognitive functions (attention: Thiebaut de Schotten et al., 2011; self recognition: Morita et al., 2017, 2018, 2020). Although we can only speculate at this point, it will be intriguing to investigate whether the heterogeneity in age dependency of the SLF microstructure explains the heterogeneous age-dependent profiles of these functions.

### 4.2. Right lateralization of the SLF III

Previous dMRI studies have reported that the right SLF III has a larger volume than the left SLF III (Budisavljevic et al., 2015, 2017; Cazzoli & Chechlacz, 2017; Hecht et al., 2015; Thiebaut de Schotten et al., 2011). We also found a significant right lateralization of tract volume estimates in SLF III (Figure 7) in adults. In addition, we found that FA measurements along SLF III are right lateralized in adolescents, adults, and seniors (Figure 7). Right lateralization of SLF III in terms of diffusivity measurements is also consistent with previous reports (Budisavljevic et al., 2017; Cazzoli & Chechlacz, 2017; Chica et al., 2018; Ioannucci et al., 2020). The replication of these findings in this independent dataset is encouraging, given the importance of the generalization of findings in neuroimaging (Lerma-Usabiaga et al., 2019).

There are two alternative hypotheses for the age dependency of SLF III lateralization. One hypothesis is that it is established in an early developmental stage and remains stable throughout the lifespan (Budisavljevic et al., 2015). An alternative hypothesis is that it emerges during relatively later developmental stage (e.g., adolescents). Consistent with the second hypothesis, a previous developmental fMRI study reported that the activation profiles of cortical regions connected by SLF III that are induced by a proprioceptive illusory task are symmetric in children but become right lateralized during adolescence, suggesting that some aspects of lateralized cortical functions mediated by SLF III emerge during development (Naito et al., 2017). Similarly, Morita and colleagues (2018) reported that right-dominant activity in the SLF III network during a self-face recognition task emerges during adolescence, but is not present in children. In the present study, we found statistically significant evidence of SLF III lateralization in terms of tract volume in adults and in terms of FA measurements in adolescents, adults, and seniors (Figure 7). In children, we found no evidence of right lateralization either for tract volume or for FA measurements of SLF III. However, it is difficult to conclude whether such lateralization is absent in children based on the present data, since our results depended on an arbitrary selection of the statistical threshold (α = 0.004) and may have also been affected by limitations in statistical power. Therefore, based on the present data, we cannot definitively distinguish between the two hypotheses. The association between lateralized cortical functions identified in developmental fMRI studies and SLF III lateralization should be evaluated in future studies.

### 4.3. Right lateralization of the SLF II

Several previous studies have investigated the lateralization of SLF II, in addition to that of SLF III, and have reported that the volume of SLF II is symmetric when data are averaged across participants (Budisavljevic et al., 2017; Cazzoli & Chechlacz, 2017; Thiebaut de Schotten et al., 2011). In contrast, other studies have found evidence of right lateralization of SLF II in terms of anisotropy measurements of dMRI data (such as FA or hindrance modulated orientational anisotropy; Budisavljevic et al., 2017; Cazzoli & Chechlacz, 2017; Chechlacz et al., 2015a; Ioannucci et al., 2020). Our results are consistent with these previous studies; we did not find evidence regarding the lateralization of the SLF II tract volume in any age group but found evidence of SLF II lateralization of FA measurements in adolescents and adults (Figure 7). Therefore, the present results, combined with those of previous studies, suggest that although the macroscopic properties of SLF II are largely symmetric, the microstructural properties of SLF II are right lateralized, and this lateralization is already present during adolescence.

Lateralization of SLF II has also been considered to be associated with cortical functions. Previous studies have demonstrated that participants with a larger right SLF II produce a greater deviation to the left in the line bisection test, suggesting a link between SLF II lateralization and spatial attention (Cazzoli & Chechlacz, 2017; Thiebaut de Schotten et al., 2011). Budisavljevic and colleagues (2017) demonstrated that the LI of SLF II is correlated with amplitude peak acceleration during a reaching task, suggesting that SLF II lateralization associated with visuomotor processing. Therefore, understanding the age dependence and microstructural basis of SLF II lateralization is essential for understanding cortical functions associated with spatial attention or visuomotor control.

### 4.4. Left lateralization of the SLF I

We found significant evidence of left lateralization of the SLF I in terms of qR1 measurements in adults (Figure 7). Since SLF I is involved in the communication between the parietal cortex and supplementary motor area (Makris et al., 2005), we speculate that this microstructural asymmetry, presumably associated with myelin levels, is involved in motor functions. Consistent with this idea, a previous study reported that the laterality of SLF I volume is significantly different between left-handed and right-handed individuals, suggesting that SLF I lateralization is associated with hand preference (Howells et al., 2018). Since the majority of the participants analyzed in this study were right-handed, we speculate that left lateralization of qR1 along SLF I may be associated with hand preference. Our supplementary analysis including only right-handed participants also supports SLF I left lateralization of qR1 measurements in adults (Supplementary Figure 13).

### 4.5. Distinct lateralization profile between macrostructural and microstructural measurements

In our study, the degree of lateralization was not fully consistent between macrostructural (tract volume) and microstructural measurements (FA and qR1) on each SLF branch. Specifically, although there was a significant right lateralization of SLF II in terms of FA measurements, we did not find a significant lateralization of SLF II tract volume (Figure 7). We speculate that the results of tract volume and FA are dissociated because they reflect different factors determining the properties of white matter tracts. Since SLF I, II, and III are segregated by sulci (Petrides & Pandya, 1984; Schmahmann & Pandya, 2006; Thiebaut de Schotten et al., 2011), the biggest constraint for the tract volume of each branch is the white matter volume of each gyrus. Therefore, we consider that cortical folding pattern (Zilles et al., 2013) may be a major factor for determining the lateralization of the tract volume. In contrast, FA measurements are affected by microstructural properties along the tract, such as axon diameter or myelin (Assaf et al., 2019; Sampaio-Baptista & Johansen-Berg, 2017). While cortical folding and microstructural properties may not be fully independent (Ecker et al., 2016), we hypothesize that macrostructural and microstructural lateralization of SLF II may be determined by different types of neurobiological mechanisms.

### 4.6. Distinct microstructural lateralization profile between SLF I and SLF II/III

A number of previous studies have proposed that FA and qR1 measurements may reflect distinct types of microstructural properties of the white matter. For example, FA is known to be sensitive to multiple types of microstructural properties including axonal properties, crossing fibers and myelination (Assaf et al., 2019; Jones et al., 2013; Rokem et al., 2017; Sampaio-Baptista & Johansen-Berg, 2017; Thomason & Thompson, 2011; Wandell & Le, 2017). In contrast, qR1 has been considered to be relatively specific to myelin levels in the cerebral white matter (Schurr et al., 2018; Stüber et al., 2014). In fact, previous studies have revealed that diffusivity measurements and qR1 measurements often provide a distinct profile along white matter tracts in relation to age dependency or disorders (Takemura et al., 2019; Yeatman et al., 2014).

Our results demonstrate that FA and qR1 measurements are sensitive to different types of lateralization in the human SLF. Specifically, while FA was sensitive to right lateralization for SLF II/III, qR1 was sensitive to left lateralization for SLF I (Figure 7). These distinct results suggest that the microstructural basis of lateralization may differ between SLF I and SLF II/III. Since the lateralization of SLF II/III was identified by FA but not by qR1, it may be explained by microstructural differences across hemispheres other than the degree of myelination. Similarly, we speculate that myelination could be a major biological cause for left lateralization of SLF I, as supported by qR1 measurements.

However, recent studies have also identified factors other than myelin, such as axon diameter (Harkins et al., 2016) or iron concentrations (Desmond et al., 2016; Stüber et al., 2014), that affect qR1 measurements. Since it is likely that the degree of correlation between qR1 and myelination varies across brain regions, future anatomical work focusing on SLF I is necessary to establish the definitive interpretation of lateralization of qR1 measurements.

## 5. Conclusions

Our analysis of dMRI and qR1 datasets from participants of varying ages provided several primary findings. First, it revealed that age dependency quantified by FA measurements is heterogeneous among the three branches of the SLF. Second, it replicated the key observation of SLF III right lateralization, as shown in previous studies (Cazzoli & Chechlacz, 2017; Thiebaut de Schotten et al., 2011), based on an independent dataset collected from participants of various ages. Third, lateralization was observed in a specific combination of MRI measurements and SLF branches (FA in SLF II/III right lateralization; qR1 in SLF I left lateralization), suggesting that the microstructural basis of lateralization may differ among SLF branches. Overall, our results provide further evidence that the human SLF is heterogeneous in terms of age dependency, microstructural properties, and lateralization. Therefore, this study opens an avenue for understanding lateralization, development and aging of cortical functions mediated by the fronto-parietal network.

## Acknowledgments

We thank Brian Wandell, Lee Michael Perry, Aviv Mezer, and Jason Yeatman for providing the MRI dataset, Michel Thiebaut de Schotten for comments on an earlier version of the manuscript, and Yusuke Sakai in support of the data analysis.

## Data availability

The data and code for replicating analyses performed in this work will be publicly available at GitHub (https://github.com/htakemur/SLFbranchesAgedependency).

