## Supplementary information for "Age dependency and lateralization in the three branches of the human superior longitudinal fasciculus"

**Affiliations:**

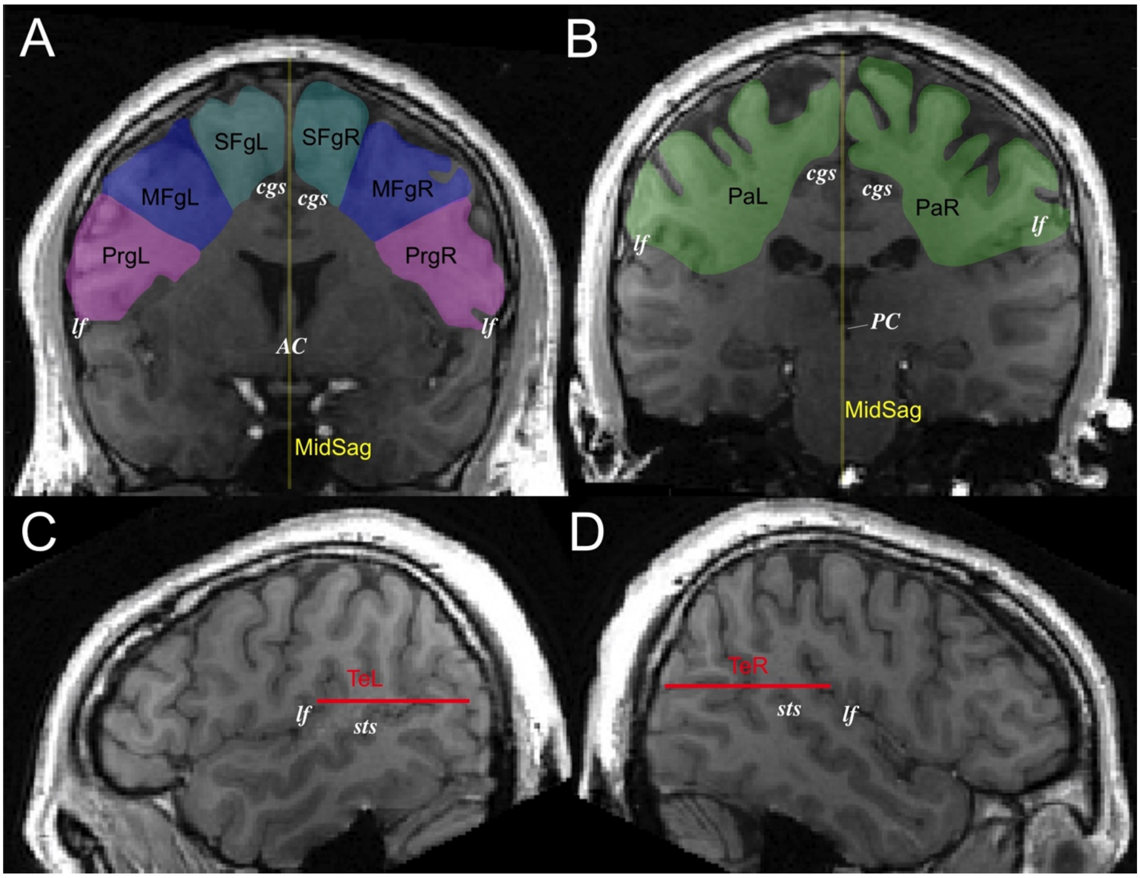


**Supplementary Figure 1.** Position of inclusion and exclusion regions of interest (ROIs) used for identifying SLF I, II, and III, overlaid on a synthetic T1-weighted image in a representative participant (P38, adult). **A.** Coronal AND ROIs, which cover the superior frontal gyrus (cyan, SFgL/SFgR), middle frontal gyrus (blue, MFgL/MFgR), and precentral gyrus (magenta, PrgL/PrgR) at the level of the anterior commissure (AC). Each of ROI was used to identify SLF I, II, and III, respectively. ROIs did not include the white matter near the cingulate gyrus, which is under the cingulate sulcus (cgs). We also defined a NOT ROI in the mid-sagittal plane (yellow, MidSag). lf: lateral fissure. **B.** Coronal AND ROIs (green, PaL/PaR) located at the level of the posterior commissure (PC). ROI covers the white matter areas superior to the lateral fissure in each hemisphere, but does not include areas near the cingulate gyrus. **C-D.** Axial NOT ROIs in the left (**C**, red, TeL) and right hemisphere (**D**, red, TeR) for excluding the arcuate fasciculus. sts: superior temporal sulcus.

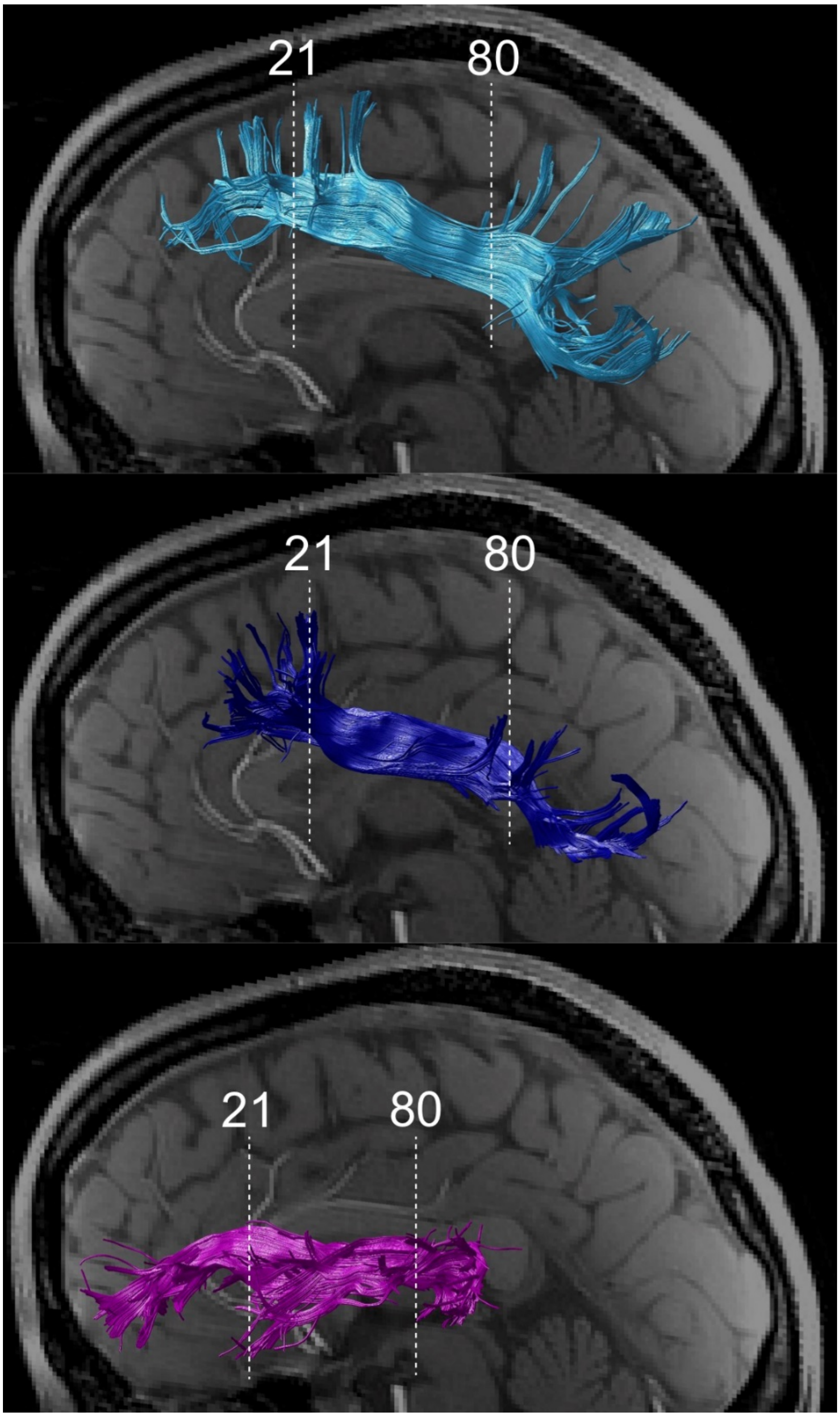


**Supplementary Figure 2.** Position of nodes of tracts included in the analyses. White dotted lines show the position of node 21 and 80 along the tract core of SLF I (cyan, top panel), SLF II (dark blue, middle panel), and SLF III (magenta, bottom panel) in the anterior-posterior coordinate of a representative participant (P18, adolescent). We performed analyses on FA and qR1 along each tract, which were averaged from node 21 and 80 in each individual hemisphere.

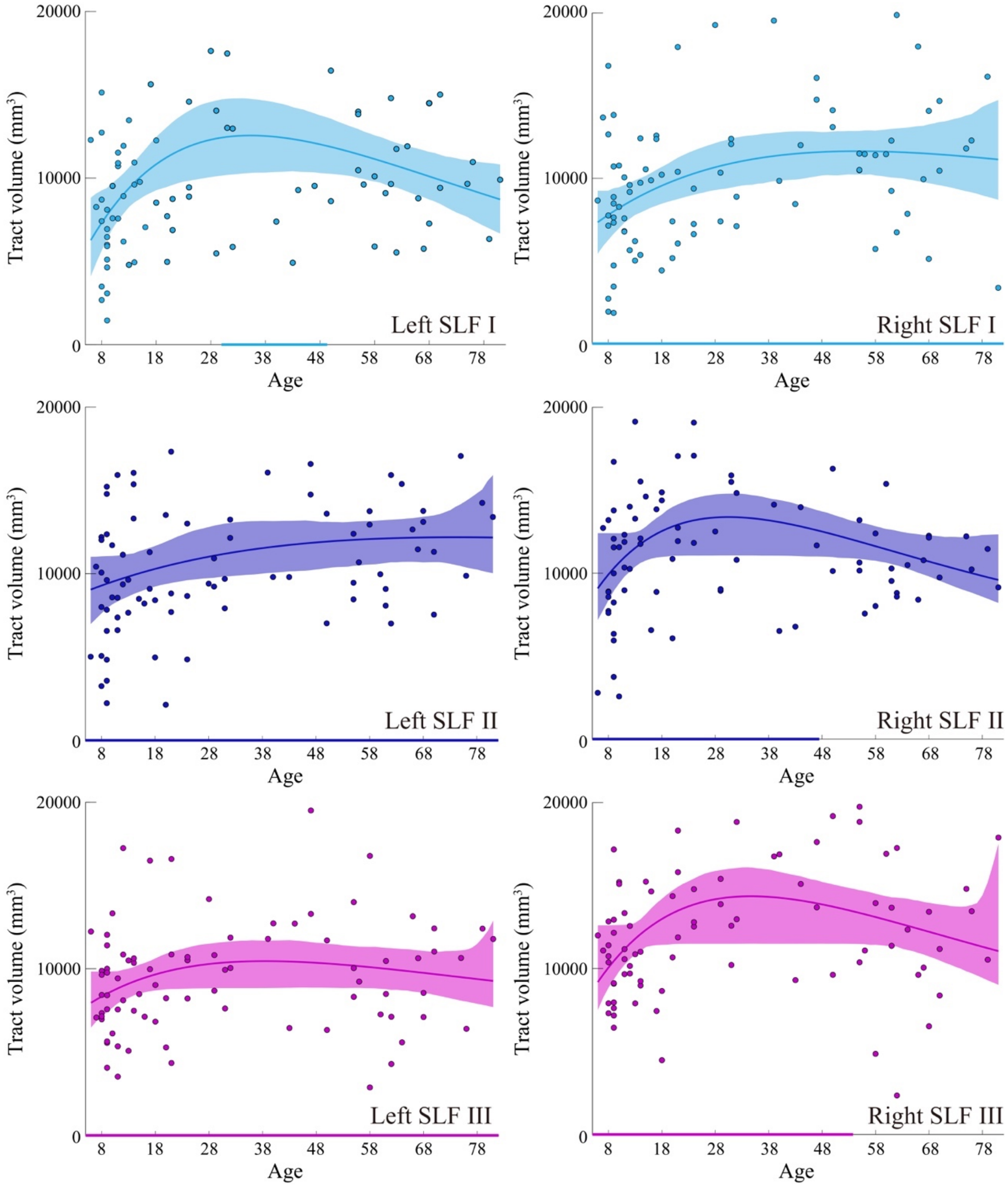


**Supplementary Figure 3.** Tract volume age-dependency curves in each SLF branch (left/right SLF I, II, and III). Each dot depicts data for an individual participant. The width of the curve denotes the 95% confidence interval around the Poisson curve fit. Colored lines at the bottom depicts the 95% confidence interval of peak age during the age-dependency curves, estimated by the bootstrapping method (Yeatman et al., 2014).

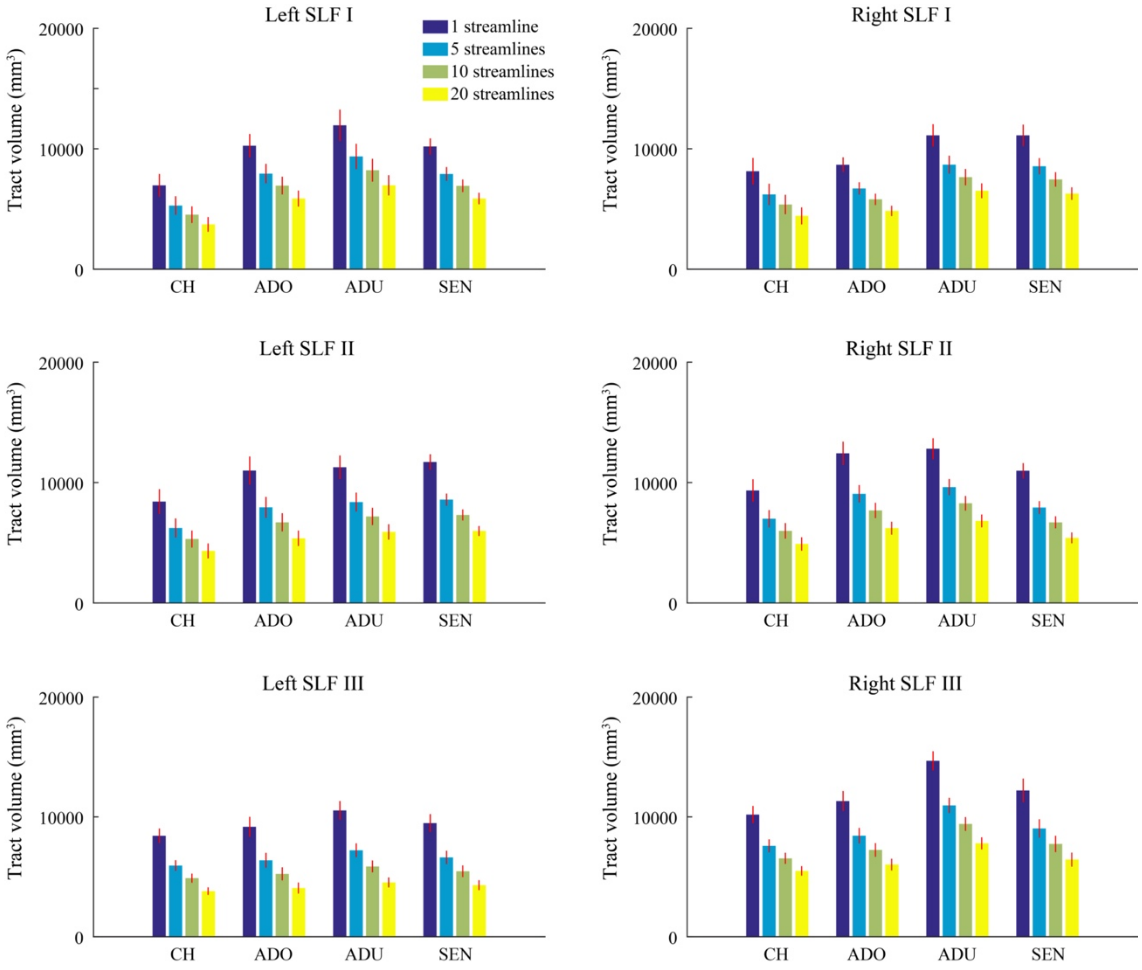


**Supplementary Figure 4.** Tract volume of the three branches of the SLF in the left and right hemisphere in each age group estimated by using four different streamline density thresholds (1, 5, 10, and 20 streamlines per voxel for dark blue, light blue, light green, and yellow). Other conventions are identical to those used in Figure 2. See Supplementary Table 1 for statistics. CH, child; ADO, adolescent; ADU, adult; SEN, senior.

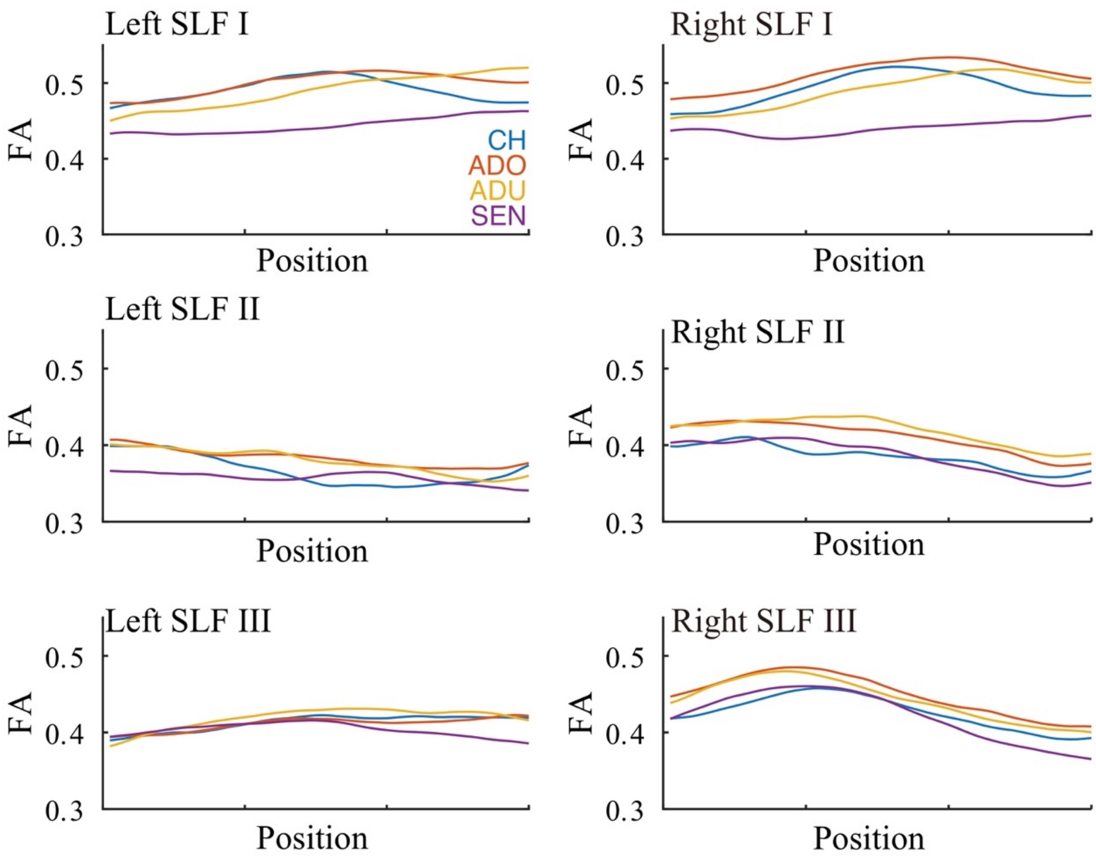


**Supplementary Figure 5.** Spatial profile of fractional anisotropy (FA) along the tract. The horizontal axis depicts spatial position along the SLF branches (left side, anterior; right side, posterior; nodes 21-80, as shown in Supplementary Figure 2). The vertical axis depicts FA. Each colored curve depicts the spatial profile of FA averaged across participants in each group (blue, child [CH]; red, adolescent [ADO]; yellow, adult [ADU]; purple, senior [SEN]).

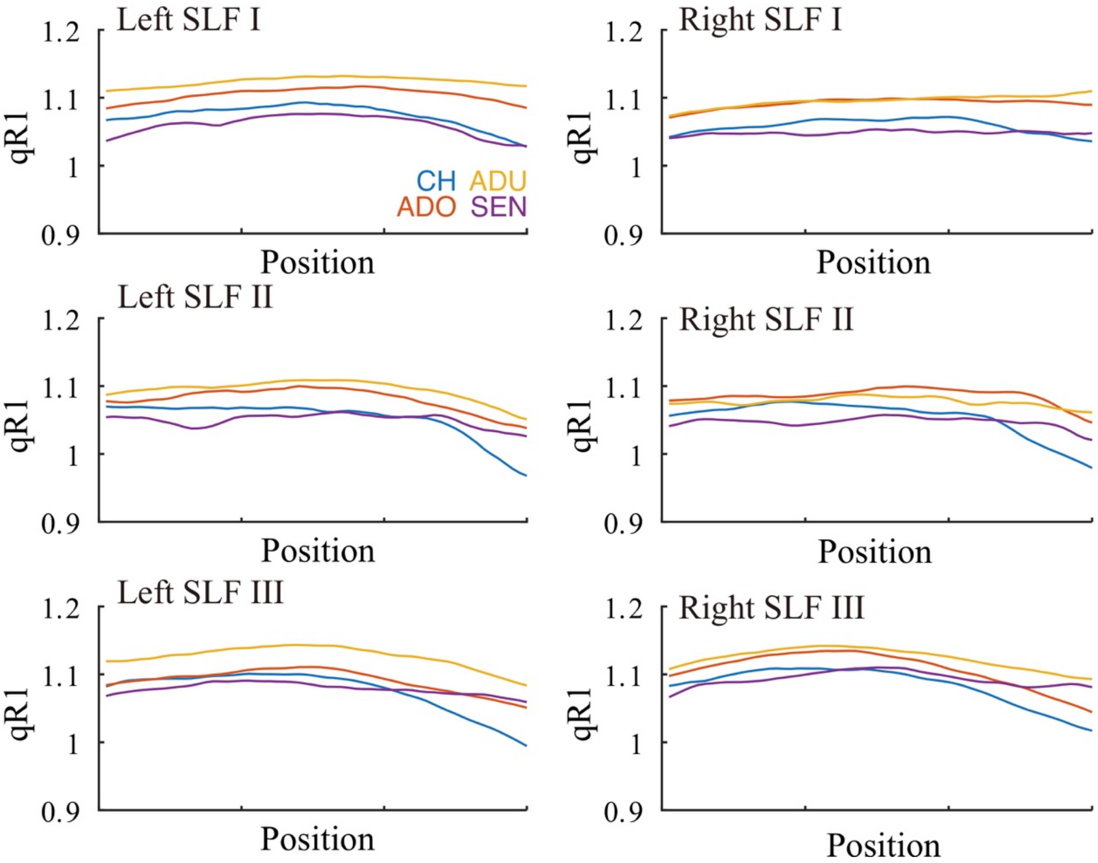


**Supplementary Figure 6.** Spatial profile of quantitative R1 (qR1) along the tract. Conventions are identical to those used in Supplementary Figure 5. CH, child; ADO, adolescent; ADU, adult; SEN, senior.

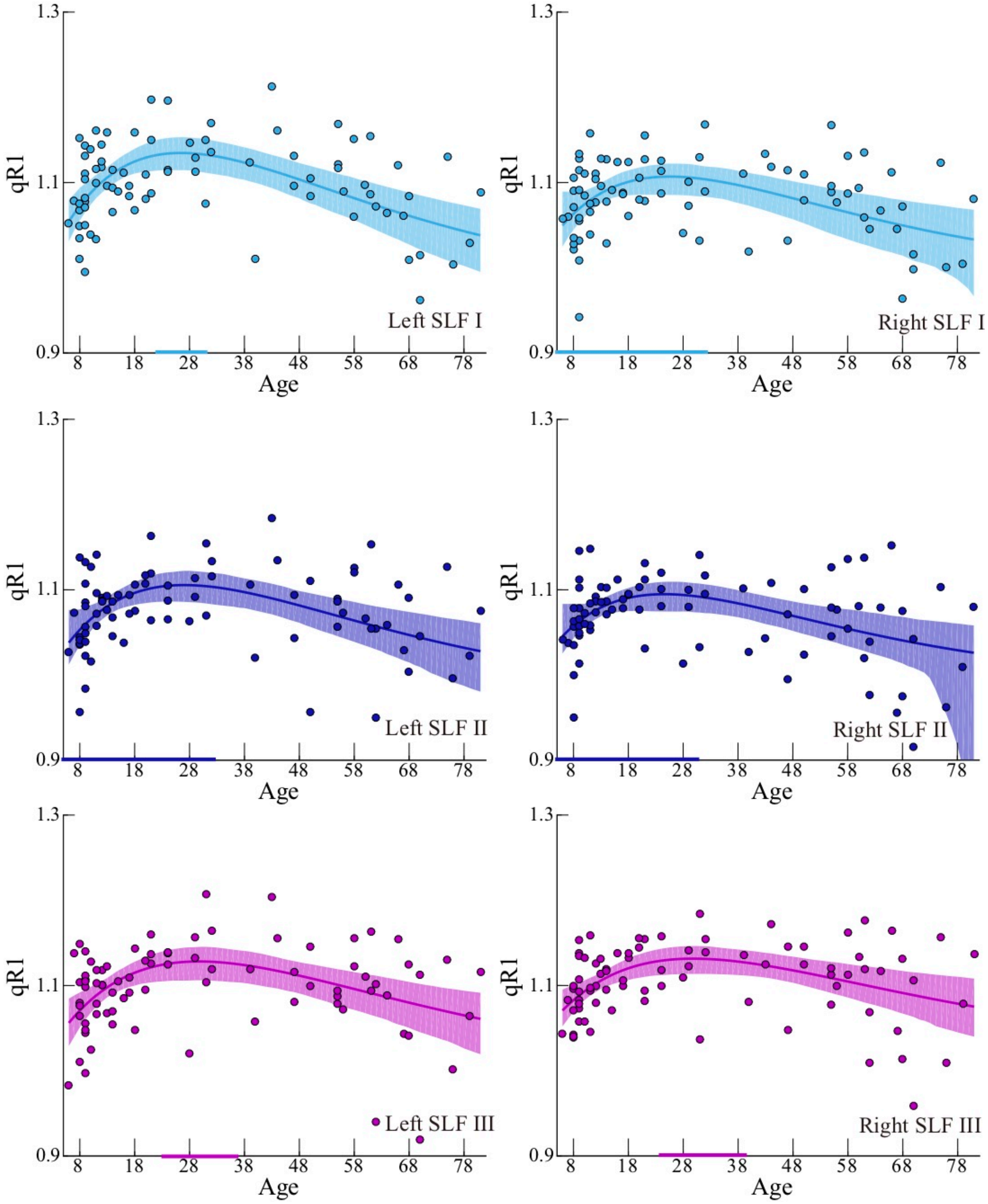


**Supplementary Figure 7.** Quantitative R1 (qR1) age-dependency curves for the SLF branches. Conventions are identical to those in Supplementary Figure 3.

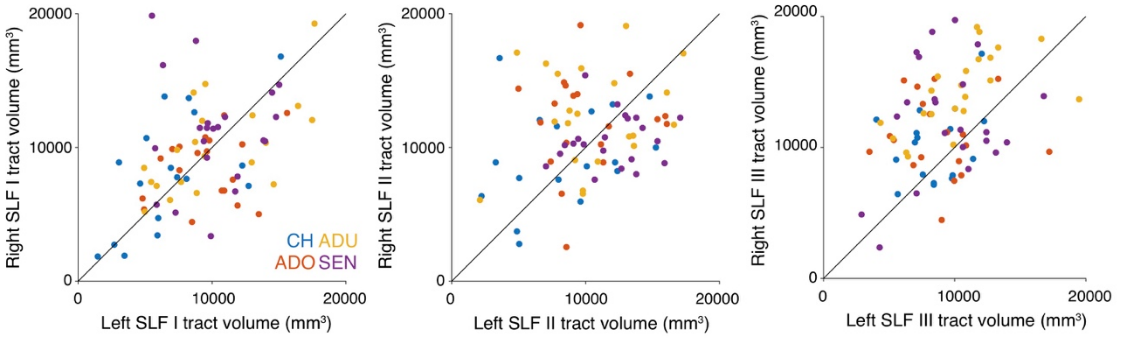


**Supplementary Figure 8.** Scatter plots comparing the tract volume between hemispheres (left, SLF I; middle, SLF II; right, SLF III). Individual dots depict data in individual participants. Color indicates age groups (blue, child [CH]; red, adolescent [ADO]; yellow, adult [ADU]; purple, senior [SEN]).

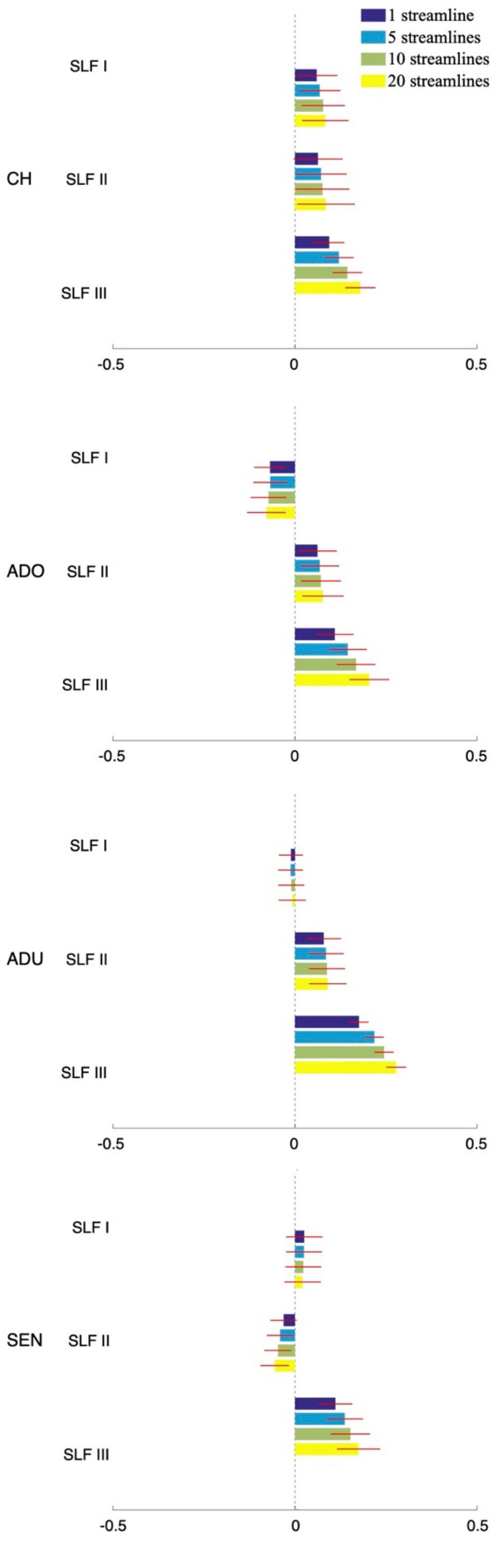


**Supplementary Figure 9.** Lateralization index of tract volume of the SLF in the left and right hemisphere in each age group estimated by using four different streamline density thresholds (1, 5, 10, and 20 streamlines per voxel for dark blue, light blue, light green, and yellow). Other conventions are identical to those used in Figure 7. CH, child; ADO, adolescent; ADU, adult; SEN, senior.

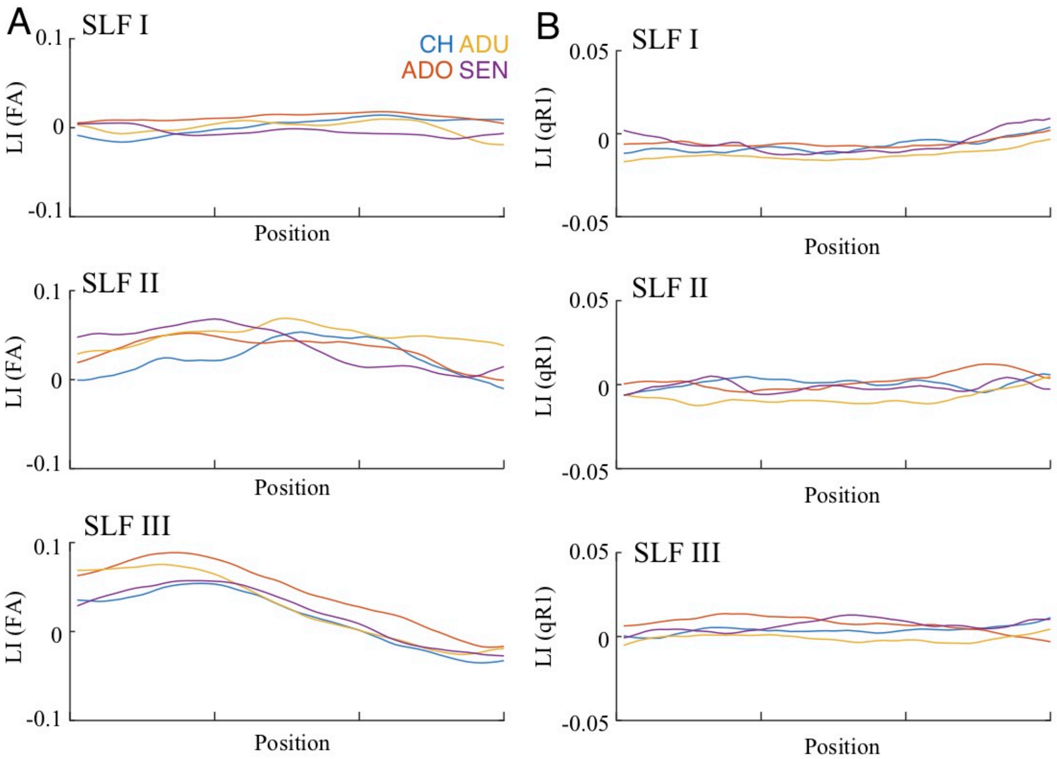


**Supplementary Figure 10.** Spatial profile of the lateralization index (LI; **A**, fractional anisotropy [FA]; **B**, quantitative R1 [qR1]). Conventions are identical to those used in Supplementary Figure 5. CH, child; ADO, adolescent; ADU, adult; SEN, senior.

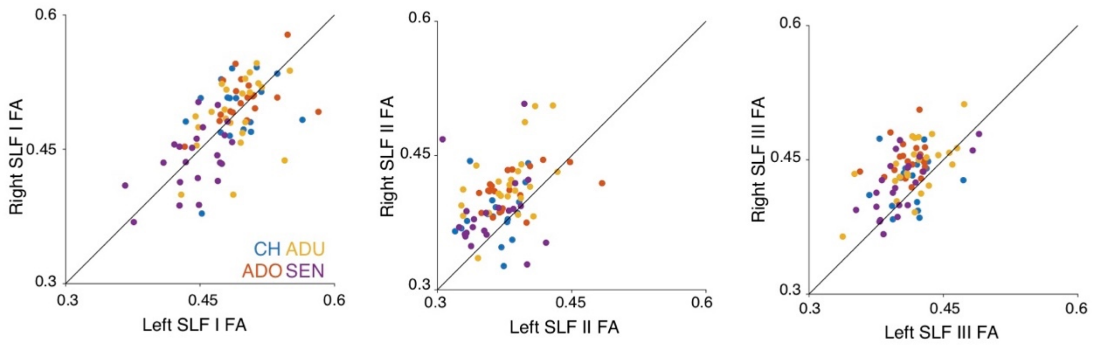


**Supplementary Figure 11.** Scatter plots comparing the fractional anisotropy (FA) between hemispheres (left, SLF I; middle, SLF II; right, SLF III). Conventions are identical to those used in Supplementary Figure 8. CH, child; ADO, adolescent; ADU, adult; SEN, senior.

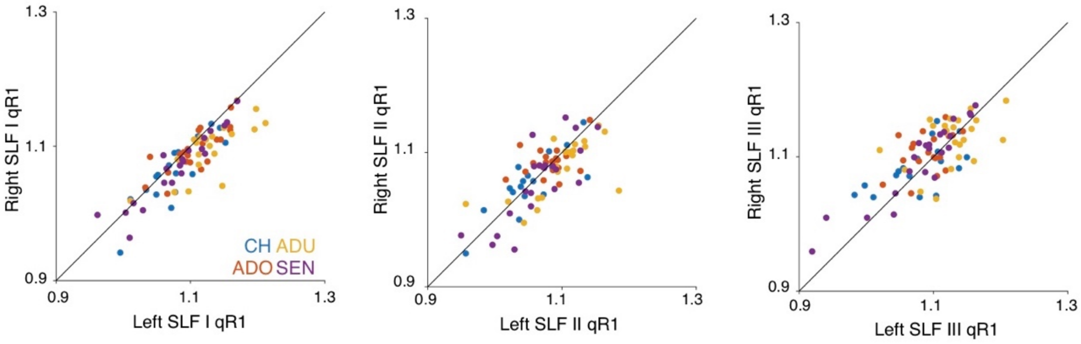


**Supplementary Figure 12.** Scatter plots comparing the quantitative R1 (qR1) between hemispheres (left, SLF I; middle, SLF II; right, SLF III). Conventions are identical to those used in Supplementary Figure 8. CH, child; ADO, adolescent; ADU, adult; SEN, senior.

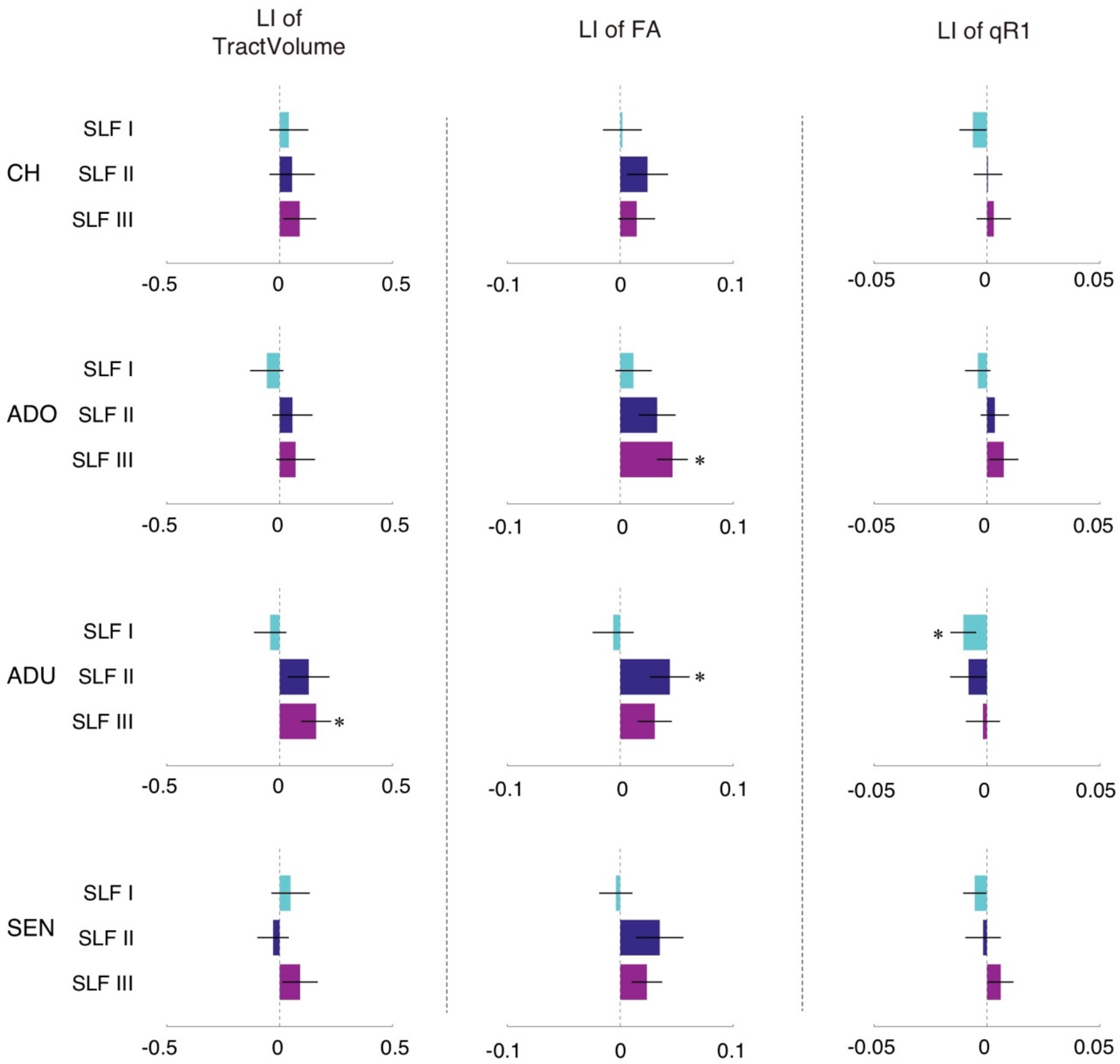


**Supplementary Figure 13.** Lateralization index (LI) in right-handers. Horizontal axis depicts the LI calculated only from right-handed participants. Conventions are identical to those in Figure 7. CH, child; ADO, adolescent; ADU, adult; SEN, senior.

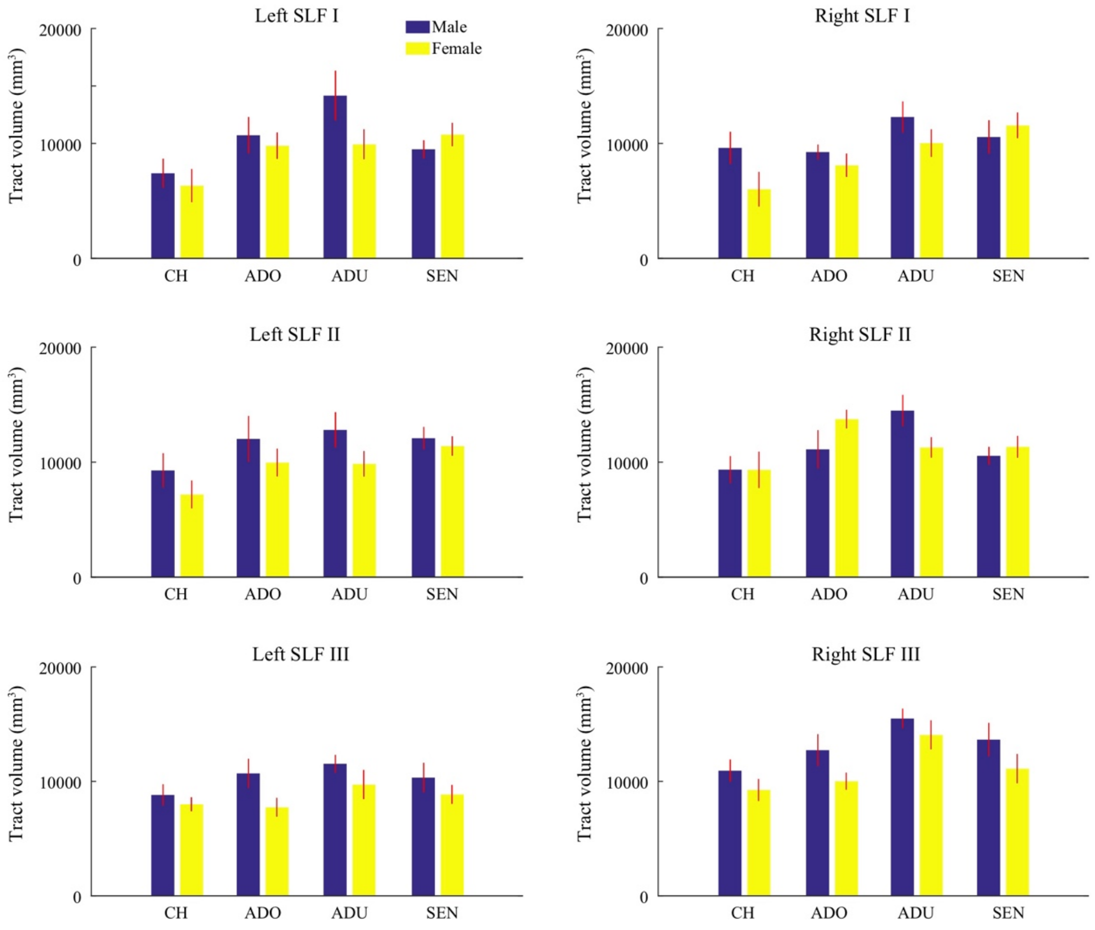


**Supplementary Figure 14.** Comparison of tract volume of the three branches of the SLF in the left and right hemisphere in each age group between male (blue) and female (yellow) participants. Other conventions are identical to those in Supplementary Figure 4. CH, child; ADO, adolescent; ADU, adult; SEN, senior.

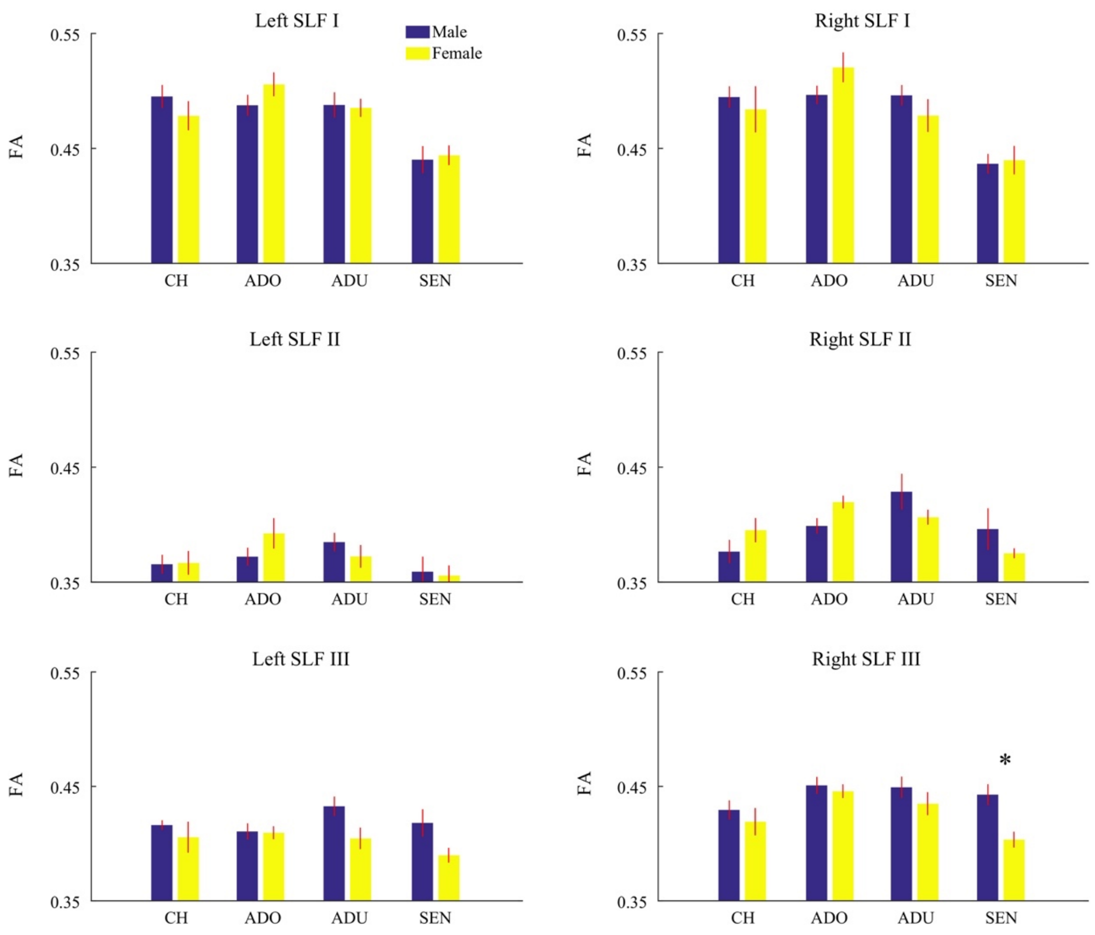


**Supplementary Figure 15.** Comparison of fractional anisotropy (FA) along three branches of the SLF in the left and right hemisphere in each age group between male (blue) and female (yellow) participants. Asterisk indicates statistically significant difference ($P < 0.002$; Bonferroni correction for 24 comparisons; see Supplementary Table 2). Other conventions are identical to those in Supplementary Figure 14. CH, child; ADO, adolescent; ADU, adult; SEN, senior.

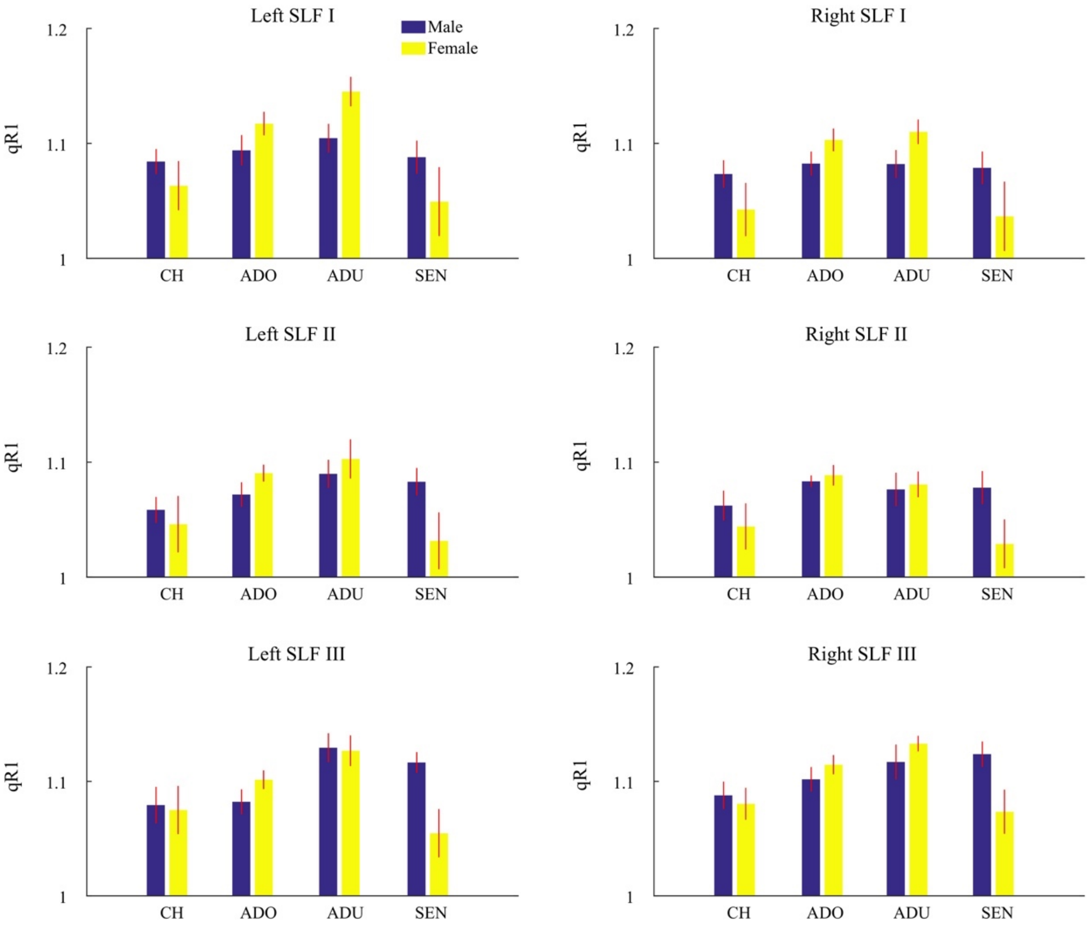


**Supplementary Figure 16.** Comparison of quantitative R1 (qR1) along three branches of the SLF in the left and right hemisphere in each age group between male (blue) and female (yellow) participants. Other conventions are identical to those used in Supplementary Figure 15. CH, child; ADO, adolescent; ADU, adult; SEN, senior.

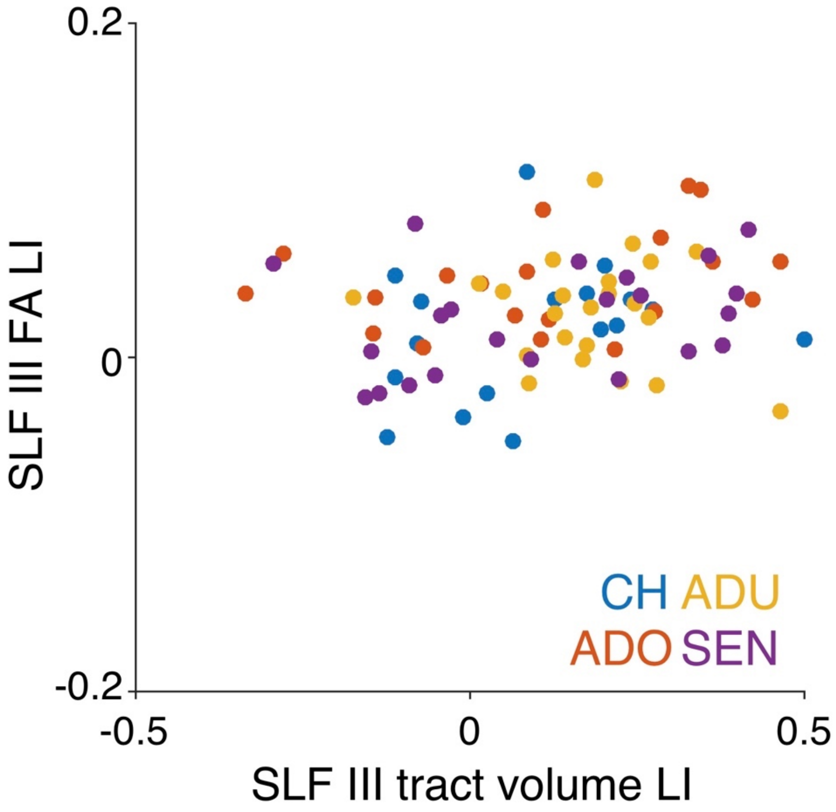


**Supplementary Figure 17.** The scatter plot comparing the laterality index (LI) of SLF III tract volume (horizontal axis) and LI of fractional anisotropy (FA) measurements (vertical axis) on the SLF III. Colored dots depict data in individual participants (blue, child [CH]; red, adolescent [ADO]; yellow, adult [ADU]; purple, senior [SEN]).

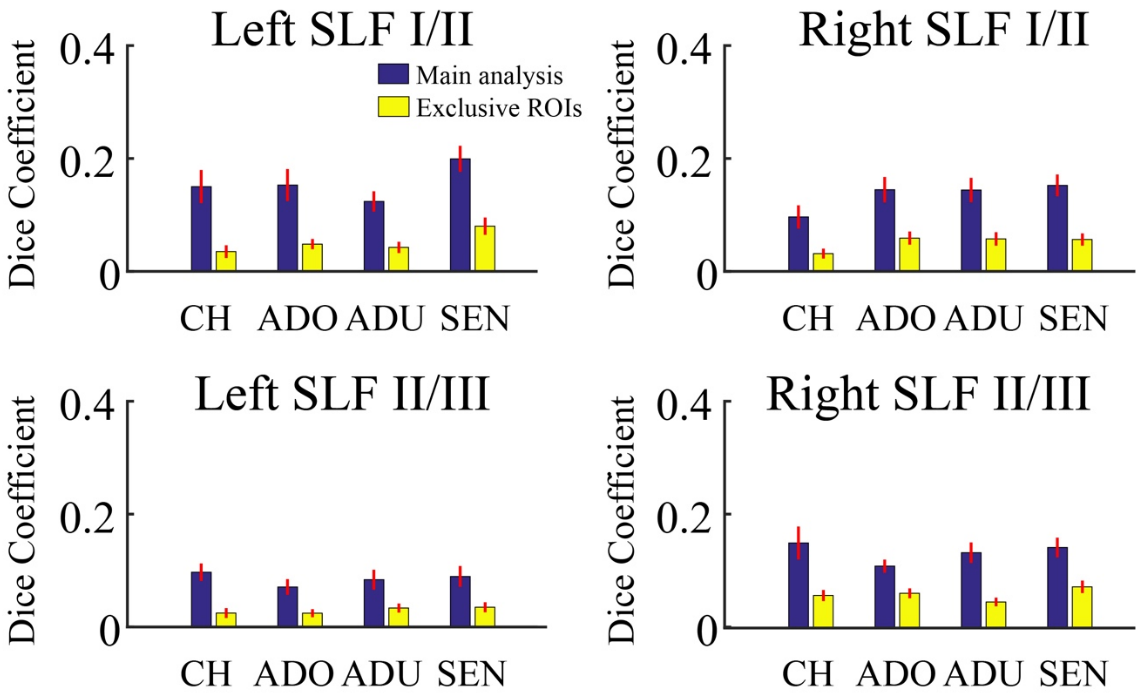


**Supplementary Figure 18.** Spatial overlap between three SLF branches. The vertical axis depicts Dice Coefficient on a proportion of overlapping voxels between SLF branches (top panels: SLF I/II, bottom panels: SLF II/III) in each age group. Blue bars depict spatial overlaps in the main analysis, while yellow bars depict spatial overlaps in supplementary analysis using exclusive ROIs (see Material and Methods). The error bars indicate ±1 s.e.m. across all participants. CH, child; ADO, adolescent; ADU, adult; SEN, senior.

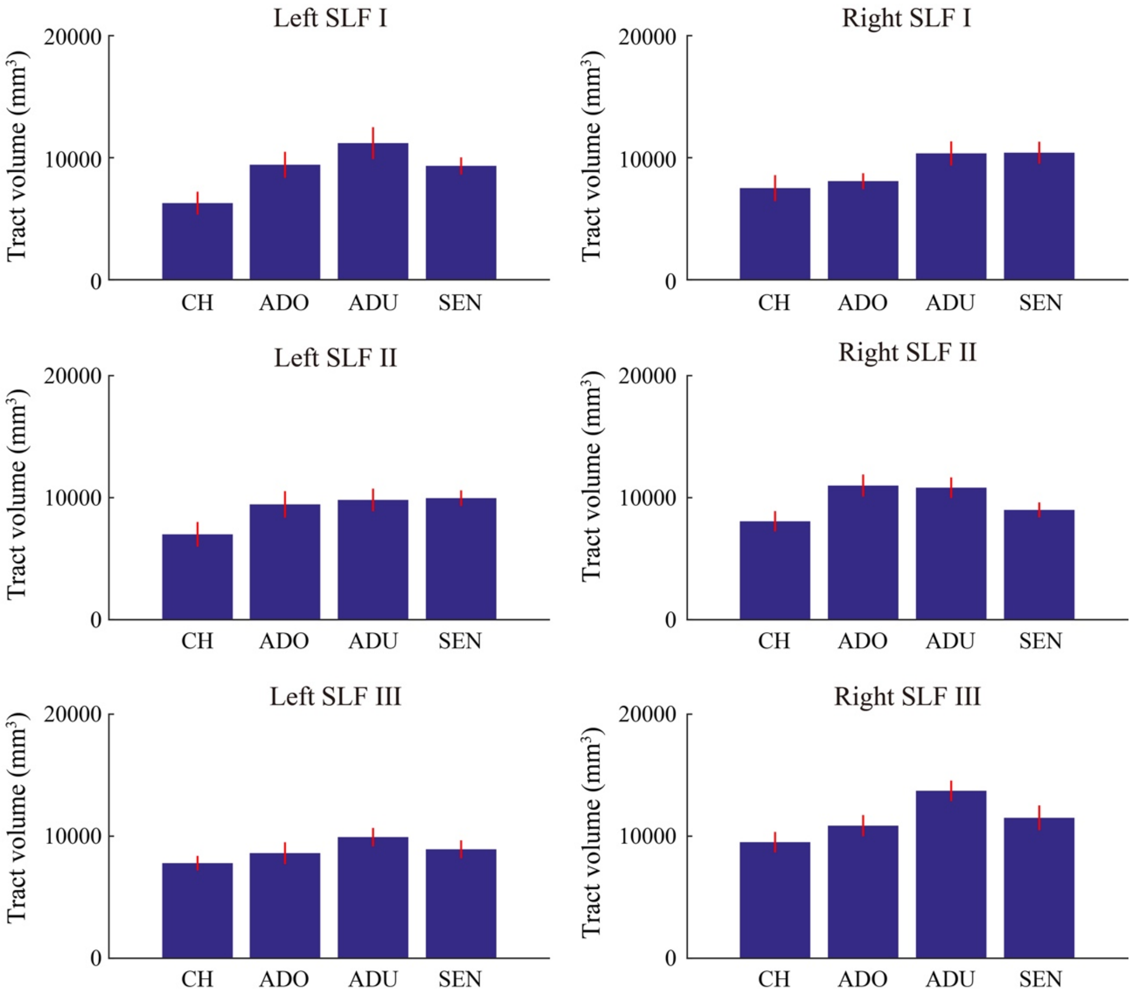


**Supplementary Figure 19.** Estimated tract volume of the three branches of the SLF identified by using exclusive ROIs (see Material and Methods). Conventions are identical to those used in Figure 2. CH, child; ADO, adolescent; ADU, adult; SEN, senior.

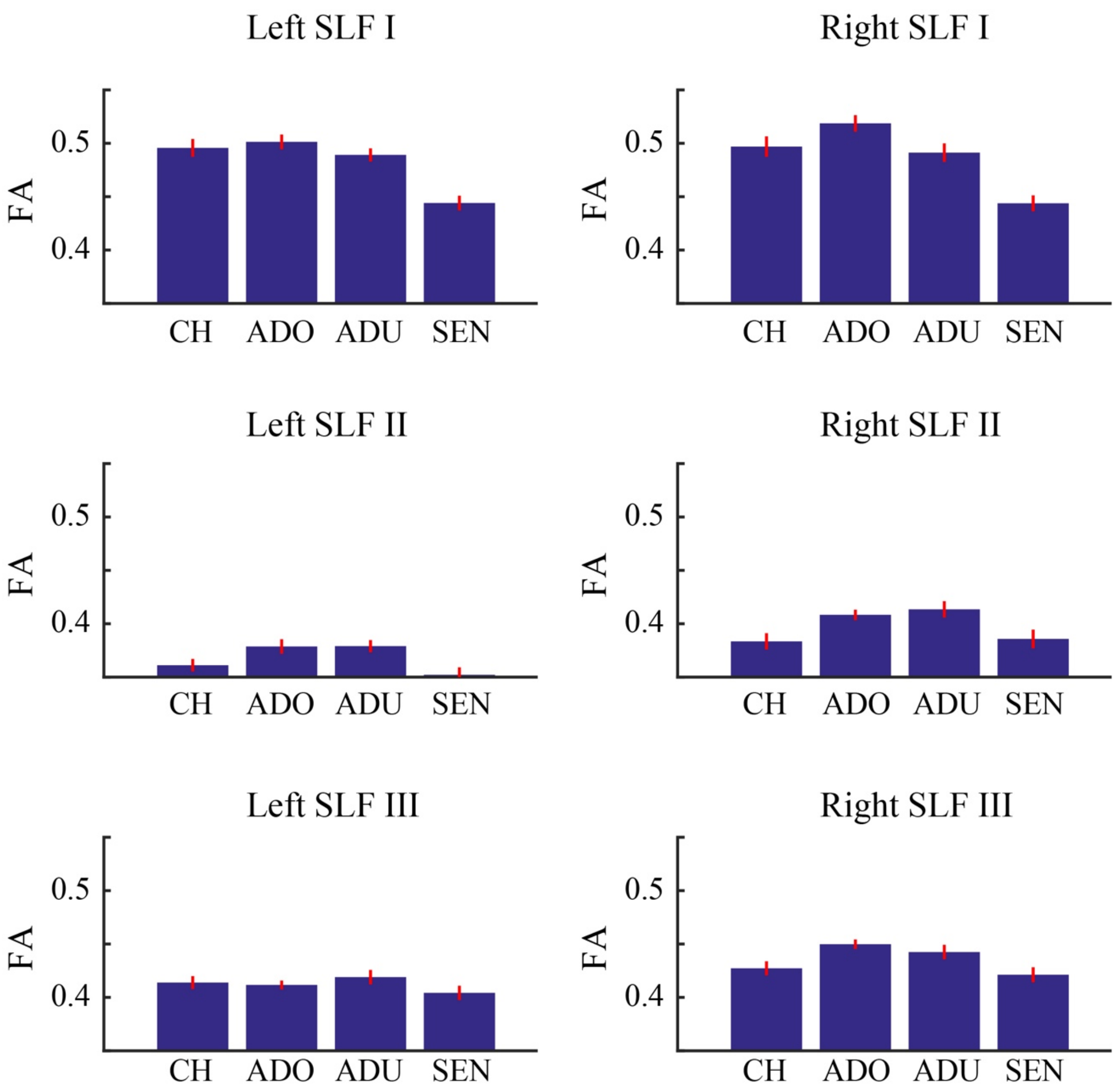


**Supplementary Figure 20.** Estimated fractional anisotropy (FA) of the three branches of the SLF identified by using exclusive ROIs. Conventions are identical to those used in Figure 3. CH, child; ADO, adolescent; ADU, adult; SEN, senior.

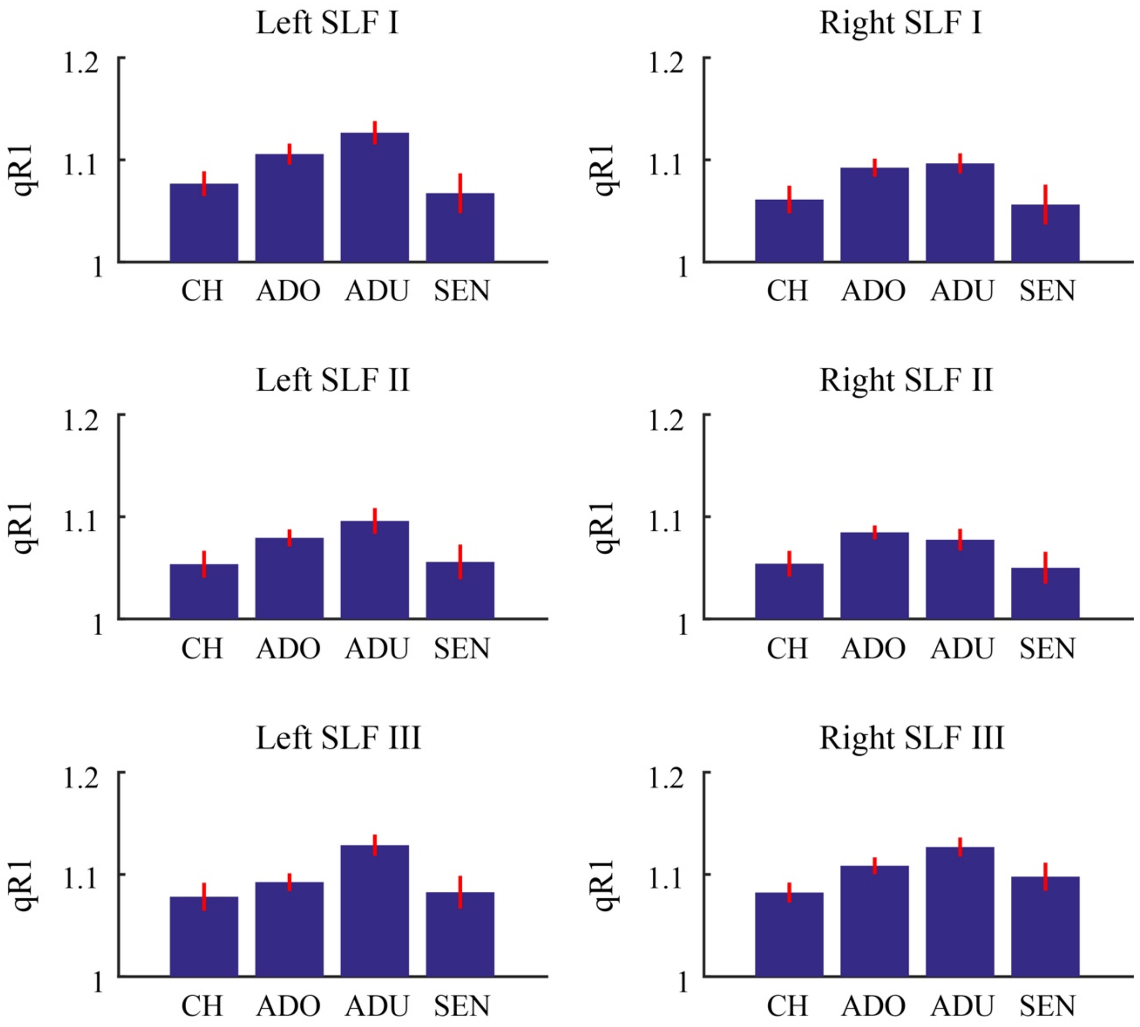


**Supplementary Figure 21.** Estimated quantitative R1 (qR1) of the three branches of the SLF identified by using exclusive ROIs. Conventions are identical to those used in Figure 6. CH, child; ADO, adolescent; ADU, adult; SEN, senior.

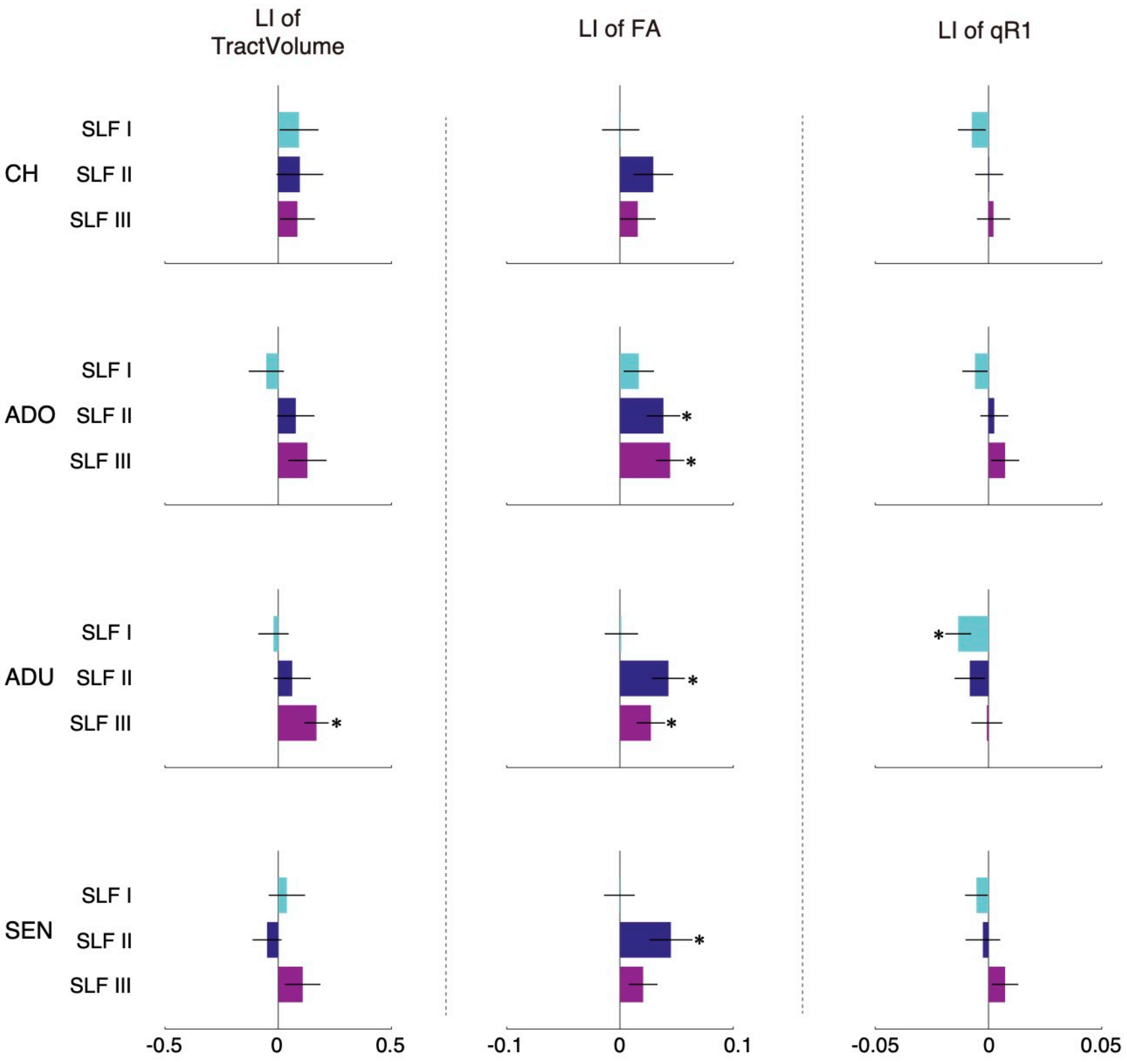


**Supplementary Figure 22.** Lateralization index of tract volume, fractional anisotropy (FA), and quantitative R1 (qR1) in the SLF I, II, and III identified by using exclusive ROIs. Conventions are identical to those used in Figure 7. CH, child; ADO, adolescent; ADU, adult; SEN, senior.

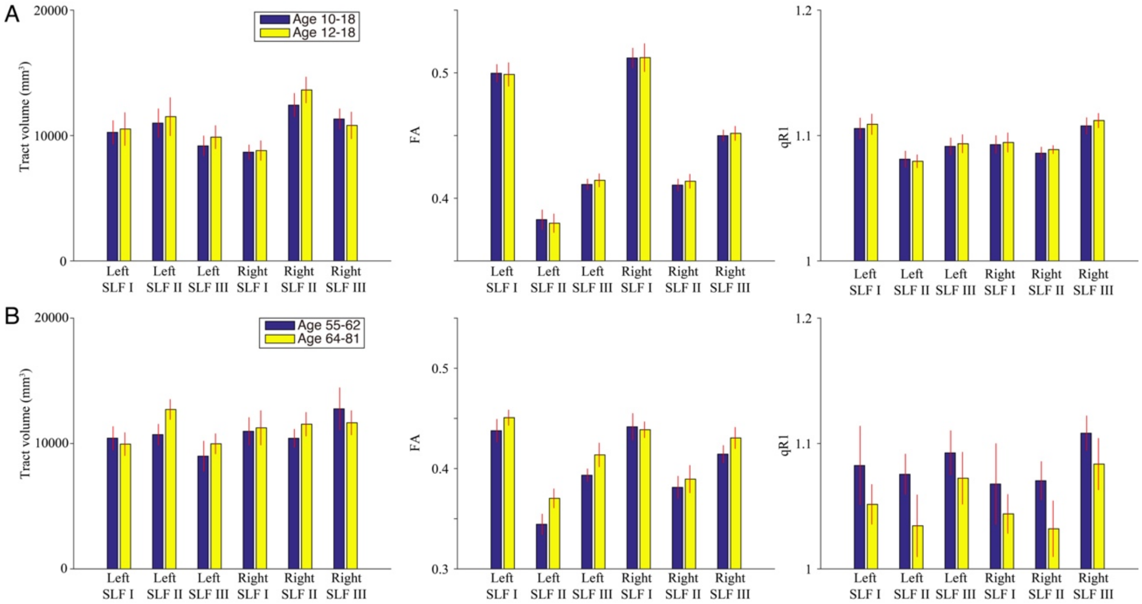


**Supplementary Figure 23.** Comparisons of results across different age group definitions. **A.** Tract volume (left), fractional anisotropy (FA; middle), and quantitative R1 (qR1; right) of each SLF branch in adolescents. The colored bars depict results on a different age group definition (blue, age 10-18, identical to the main analysis; yellow, age 12-18). **B.** Tract volume (left), FA (middle), and qR1 (right) of each SLF branch in subgroups of senior participants (blue, age 55-62; yellow, age 64-81). The error bars indicate ±1 s.e.m. across all participants.

**Supplementary Table 1. The two-way ANOVA table for tract volume when varying streamline density threshold.** Bold Italic texts indicate statistically significant effects ($P$ < 0.05).

| *Two-way ANOVA for tract volume (threshold = 1 streamline)* | | |
|---|---|---|
| | F-value | P-value |
| ***age group*** | ***$F^{3,468}$ = 16.20*** | ***P < .001*** |
| ***tract*** | ***$F^{5,468}$ = 5.99*** | ***P < .001*** |
| age group * tract | $F^{15,468}$ = 0.90 | P = 0.57 |
| *Two-way ANOVA for tract volume (threshold = 5 streamlines)* | | |
| | F-value | P-value |
| ***age group*** | ***$F^{3,468}$ = 15.80*** | ***P < .001*** |
| ***tract*** | ***$F^{5,468}$ = 6.32*** | ***P < .001*** |
| age group * tract | $F^{15,468}$ = 0.93 | P = 0.53 |
| *Two-way ANOVA for tract volume (threshold = 10 streamlines)* | | |
| | F-value | P-value |
| ***age group*** | ***$F^{3,468}$ = 15.45*** | ***P < .001*** |
| ***tract*** | ***$F^{5,468}$ = 7.11*** | ***P < .001*** |
| age group * tract | $F^{15,468}$ = 0.95 | P = 0.51 |
| *Two-way ANOVA for tract volume (threshold = 20 streamlines)* | | |
| | F-value | P-value |
| ***age group*** | ***$F^{3,468}$ = 14.82*** | ***P < .001*** |
| ***tract*** | ***$F^{5,468}$ = 8.44*** | ***P < .001*** |
| age group * tract | $F^{15,468}$ = 0.96 | P = 0.50 |

**Supplementary Table 2. Two-sample t-test comparing male and female participants.** Bold Italic texts indicate statistically significant effects ($P < 0.002$; Bonferroni corrected for 24 comparisons). CH, child; ADO, adolescent; ADU, adult; SEN, senior.

| | CH | ADO | ADU | SEN |
|---|---|---|---|---|
| *Tract Volume* | | | | |
| Left SLF I | $t_{15} = 0.57, P = 0.58$ | $t_{18} = 0.48, P = 0.64$ | $t_{21} = 1.75, P = 0.09$ | $t_{20} = -1.01, P = 0.32$ |
| Left SLF II | $t_{15} = 1.04, P = 0.31$ | $t_{18} = 0.90, P = 0.38$ | $t_{21} = 1.62, P = 0.12$ | $t_{20} = 0.56, P = 0.58$ |
| Left SLF III | $t_{15} = 0.69, P = 0.50$ | $t_{18} = 2.03, P = 0.06$ | $t_{21} = 1.23, P = 0.23$ | $t_{20} = 1.03, P = 0.32$ |
| Right SLF I | $t_{15} = 1.77, P = 0.10$ | $t_{18} = 1.00, P = 0.33$ | $t_{21} = 1.30, P = 0.21$ | $t_{20} = -0.58, P = 0.57$ |
| Right SLF II | $t_{15} = 0.00, P = 0.99$ | $t_{18} = -1.46, P = 0.16$ | $t_{21} = 2.07, P = 0.05$ | $t_{20} = -0.65, P = 0.52$ |
| Right SLF III | $t_{15} = 1.24, P = 0.23$ | $t_{18} = 1.78, P = 0.09$ | $t_{21} = 0.95, P = 0.35$ | $t_{20} = 1.35, P = 0.19$ |
| *FA* | | | | |
| Left SLF I | $t_{15} = 1.08, P = 0.30$ | $t_{18} = -1.38, P = 0.18$ | $t_{21} = 0.20, P = 0.85$ | $t_{20} = -0.28, P = 0.78$ |
| Left SLF II | $t_{15} = -0.10, P = 0.92$ | $t_{18} = -1.37, P = 0.19$ | $t_{21} = 1.00, P = 0.33$ | $t_{20} = 0.23, P = 0.82$ |
| Left SLF III | $t_{15} = 0.90, P = 0.38$ | $t_{18} = 0.13, P = 0.90$ | $t_{21} = 2.31, P = 0.03$ | $t_{20} = 2.29, P = 0.03$ |
| Right SLF I | $t_{15} = 0.55, P = 0.59$ | $t_{18} = -1.63, P = 0.12$ | $t_{21} = 1.06, P = 0.30$ | $t_{20} = -0.21, P = 0.84$ |
| Right SLF II | $t_{15} = -1.30, P = 0.21$ | $t_{18} = -2.54, P = 0.02$ | $t_{21} = 1.40, P = 0.18$ | $t_{20} = 1.27, P = 0.22$ |
| Right SLF III | $t_{15} = 0.77, P = 0.46$ | $t_{18} = 0.56, P = 0.58$ | $t_{21} = 1.09, P = 0.29$ | ***$t_{20} = 3.70, P = 0.001$*** |
| *qR1* | | | | |
| Left SLF I | $t_{15} = 0.98, P = 0.34$ | $t_{18} = -1.44, P = 0.17$ | $t_{21} = -2.34, P = 0.03$ | $t_{20} = 1.11, P = 0.28$ |
| Left SLF II | $t_{15} = 0.53, P = 0.61$ | $t_{18} = -1.51, P = 0.15$ | $t_{21} = -0.63, P = 0.54$ | $t_{20} = 1.79, P = 0.09$ |
| Left SLF III | $t_{15} = 0.17, P = 0.87$ | $t_{18} = -1.49, P = 0.15$ | $t_{21} = 0.15, P = 0.89$ | $t_{20} = 2.57, P = 0.02$ |
| Right SLF I | $t_{15} = 1.33, P = 0.20$ | $t_{18} = -1.50, P = 0.15$ | $t_{21} = -1.81, P = 0.08$ | $t_{20} = 1.21, P = 0.24$ |
| Right SLF II | $t_{15} = 0.82, P = 0.43$ | $t_{18} = -0.55, P = 0.59$ | $t_{21} = -0.25, P = 0.80$ | $t_{20} = 1.87, P = 0.08$ |
| Right SLF III | $t_{15} = 0.41, P = 0.69$ | $t_{18} = -0.99, P = 0.34$ | $t_{21} = -1.03, P = 0.32$ | $t_{20} = 2.20, P = 0.04$ |

**Supplementary Table 3. Inter-individual correlation of Lateralization Index (LI) across measurements.**

| | LI of SLF II FA | LI of SLF III FA | LI of SLF I qR1 |
|---|---|---|---|
| LI of SLF III volume | R = 0.12<br>P = 0.29 | R = 0.20<br>P = 0.07 | R = -0.04<br>P = 0.72 |
| LI of SLF II FA | | R = -0.04<br>P = 0.74 | R = -0.16<br>P = 0.16 |
| LI of SLF III FA | | | R = -0.07<br>P = 0.55 |

**Supplementary Table 4. The two-way ANOVA table for tract volume, fractional anisotropy (FA), and quantitative R1 (qR1) of SLF branches identified by using exclusive ROIs.** Bold Italic texts indicate statistically significant effects (*P* < 0.05).

| | | |
|---|---|---|
| *Two-way ANOVA for tract volume* | | |
| | F-value | P-value |
| ***age group*** | ***$F^{3,468}$ = 15.44*** | ***P < .001*** |
| ***tract*** | ***$F^{5,468}$ = 5.50*** | ***P < .001*** |
| age group * tract | $F^{15,468}$ = 0.94 | P = 0.52 |
| *Two-way ANOVA for FA* | | |
| ***age group*** | ***$F^{3,468}$ = 32.77*** | ***P < .001*** |
| ***tract*** | ***$F^{5,468}$ = 183.77*** | ***P < .001*** |
| ***age group * tract*** | ***$F^{15,468}$ = 3.18*** | ***P < .001*** |
| *Two-way ANOVA for qR1* | | |
| ***age group*** | ***$F^{3,468}$ = 20.30*** | ***P < .001*** |
| ***tract*** | ***$F^{5,468}$ = 7.36*** | ***P < .001*** |
| age group * tract | $F^{15,468}$ = 0.53 | P = 0.93 |

**Supplementary Table 5. Paired t-test results on hemispheric difference of tract volume, fractional anisotropy (FA), and quantitative R1 (qR1) along SLF branches identified by using exclusive ROIs.** Bold Italic texts indicate statistically significant effects ($P$ < 0.004). CH, child; ADO, adolescent; ADU, adult; SEN, senior.

| | tract volume | FA | qR1 |
|---|---|---|---|
| *CH* | | | |
| SLF I | d’ = 0.39; P = 0.13 | d’ = 0.03; P = 0.91 | d’ = -0.56; P = 0.03 |
| SLF II | d’ = 0.27; P = 0.29 | d’ = 0.62; P = 0.02 | d’ = 0.02; P = 0.94 |
| SLF III | d’ = 0.54; P = 0.04 | d’ = 0.39; P = 0.13 | d’ = 0.11; P = 0.66 |
| *ADO* | | | |
| SLF I | d’ = -0.33; P = 0.16 | d’ = 0.48; P = 0.04 | d’ = -0.52; P = 0.03 |
| SLF II | d’ = 0.34; P = 0.14 | ***d’ = 0.91; P = 0.0007*** | d’ = 0.17; P = 0.45 |
| SLF III | d’ = 0.52; P = 0.03 | ***d’ = 1.50; P < 0.0001*** | d’ = 0.50; P = 0.04 |
| *ADU* | | | |
| SLF I | d’ = -0.22; P = 0.29 | d’ = 0.05; P = 0.81 | ***d’ = -1.00; P = 0.0001*** |
| SLF II | d’ = 0.21; P = 0.33 | ***d’ = 1.04; P = 0.0001*** | d’ = -0.47; P = 0.04 |
| SLF III | ***d’ = 1.35; P < 0.0001*** | ***d’ = 0.86; P = 0.0005*** | d’ = 0.04; P = 0.84 |
| *SEN* | | | |
| SLF I | d’ = 0.22; P = 0.31 | d’ = -0.01; P = 0.97 | d’ = -0.55; P = 0.02 |
| SLF II | d’ = -0.27; P = 0.22 | ***d’ = 0.70; P = 0.004*** | d’ = -0.13; P = 0.55 |
| SLF III | d’ = 0.55; P = 0.02 | d’ = 0.63; P = 0.01 | d’ = 0.56; P = 0.02 |
